# Drag racers and long haulers: the design space of myosin-V-like motor proteins

**DOI:** 10.64898/2026.10.25.761746

**Authors:** Joshua B. Holmes, Michael Hinczewski

## Abstract

The ubiquitous myosin class-V molecular motors transport cellular cargo long distances at roughly 0.5 *µ*m/s. Plant-specific myosin class-XI motors share nearly identical structural features with myosin-V, yet display velocities from 0.25 to 70 *µ*m/s, the fastest transport motor in nature. Empirical observations suggest an inverse relationship between run length and velocity. These observations raise a central question: how can a single motor design support both highly processive “long-hauler” and extremely rapid “drag-racer” motors? We generalize a polymer-based model of force-dependent myosin-V dynamics to encompass the broader design space. The model quantitatively captures experimental data from fast myosin-XI and yields general analytical expressions for key observables. Notably, we derive a Pareto frontier that defines the trade-off between velocity and run distance, as well as the theoretical maximum velocity achievable by such motors. Our results establish the biological limits of natural and synthetic myosin-V-like motor design.

## Introduction

Within every eukaryotic cell, thousands of myosin molecular motor proteins convert chemical energy into mechanical work by hydrolyzing ATP, powering movement along actin filaments^1^. Of the 79 genetically distinct classes of myosin motors, a significant portion traffic vesicles, mRNA, and organelles around the cell^2^. Myosin-V motors are among the best studied and understood of these intracellular transporters^3^. Myosin-V motors leverage the ATPase cycle and a dimeric structure to travel long distances along actin in 74 nm steps via a hand-over-hand mechanism^3, 4^. At the structural level, each monomer consists of a cargo binding domain connected to an extended rod region that ultimately forms a coiled coil with the second monomer^5^. Further down each monomer lies a 35 nm lever arm region that binds six calmodulin or calmodulin-like light chains in the presence of calcium ions. These light chains link end-to-end to significantly increase the lever arm’s rigidity^6^. Finally, an actin-binding head that mediates the ATPase cycle resides at the end of each monomer^7^. Since both heads must bind facing actin’s barbed end, one head becomes the trailing head while the other becomes the leading head^3^.

Since the state of the ATPase cycle determines whether a head is attached to actin, uncoordinated cycling between the two heads would cause either head to detach at random and make for a highly inefficient motor. Elegant experimental studies showed that myosin-V motors coordinate their two ATPase cycles via a gating mechanism where reverse strain on the leading lever arm slows the chemical transitions needed for actin release when both heads are bound^8, 9^. Thus, the unstrained trailing head detaches nearly ten times faster^8^. In the process, it releases the stored energy in the bent leading lever arm and springs the entire motor forward so that the free head can more easily find the forward actin-binding site^7^. Accompanying the extensive experimental understanding of myosin-V motors has been a plethora of highly successful theoretical studies^10^. Integrating structural, kinetic, and dynamic data into a single microscopic description of myosin-V gives us unprecedented insight into how specific physical parameters determine myosin-V’s functional behavior.

For all the success and attention myosin-V enjoys, however, it is still just one class of transporter myosin among many others. Another particularly intriguing group of transporters belongs to the plant-specific myosin class XI. These motors help maintaining cytoplasmic streaming within higher-order plants, which can occur at high velocities between 1 and 70 *µ*m/s^11, 12^. Structurally, myosin-XI motors are remarkably similar to myosin-V motors. They host all the same major domains and share the same lever arm length^13^. Furthermore, all existing evidence suggests that myosin-XI motors also use a hand-over-hand mechanism to travel along actin^14^. In terms of dynamics, however, myosin-XI motors are in a class of their own. One member boasts the fastest velocity measured for any molecular motor in the in vitro motility assay at 70 *µ*m/s^15^. By comparison, myosin-V tops out at only 0.5 *µ*m/s^16^. Despite the existence of these myosin-XI speed demons, not all myosin-XI motors run at such high speeds. Other myosin-XI members achieve a top speed of only 0.25 *µ*m/s^14, 15, 17^.

These observations suggest that myosin-XI and myosin-V motors exist within some larger design space of myosin-V-like motors. Developing a theoretical understanding of such a design space would not only reveal the design principles of natural myosin motor design, but inform efforts to engineer active materials based on synthetic myosins^18, 19^. To investigate the plausibility of a unifying myosin-V-like design space, we generalize an existing structural model of myosin-V dynamics and show that it can fit experimental data from *Nicotiana tabacum* (*Nt*.) myosin-XI^14^. We then explore the nature of the design space of all myosin-V-like motors, finding general expressions for observable quantities like run times, distances, and velocities. Through the model we uncover a Pareto frontier that explicitly defines a tradeoff between run distance and velocity, including an absolute maximum average run velocity that any myosin-V-like motor can obtain.

## Results

### Kinetic network of myosin dynamics

Our investigation into the design space of myosin-V-like motors starts with a well-established six-state kinetic model of myosin-V dynamics on a periodic actin track (Fig. 1). Since we largely invoke the results of the previous studies that developed and defended this model^20, 21^, we limit ourselves here to a brief description of the model’s main features. The original studies give a complete experimental justification and validation^20^.

**Figure 1.**
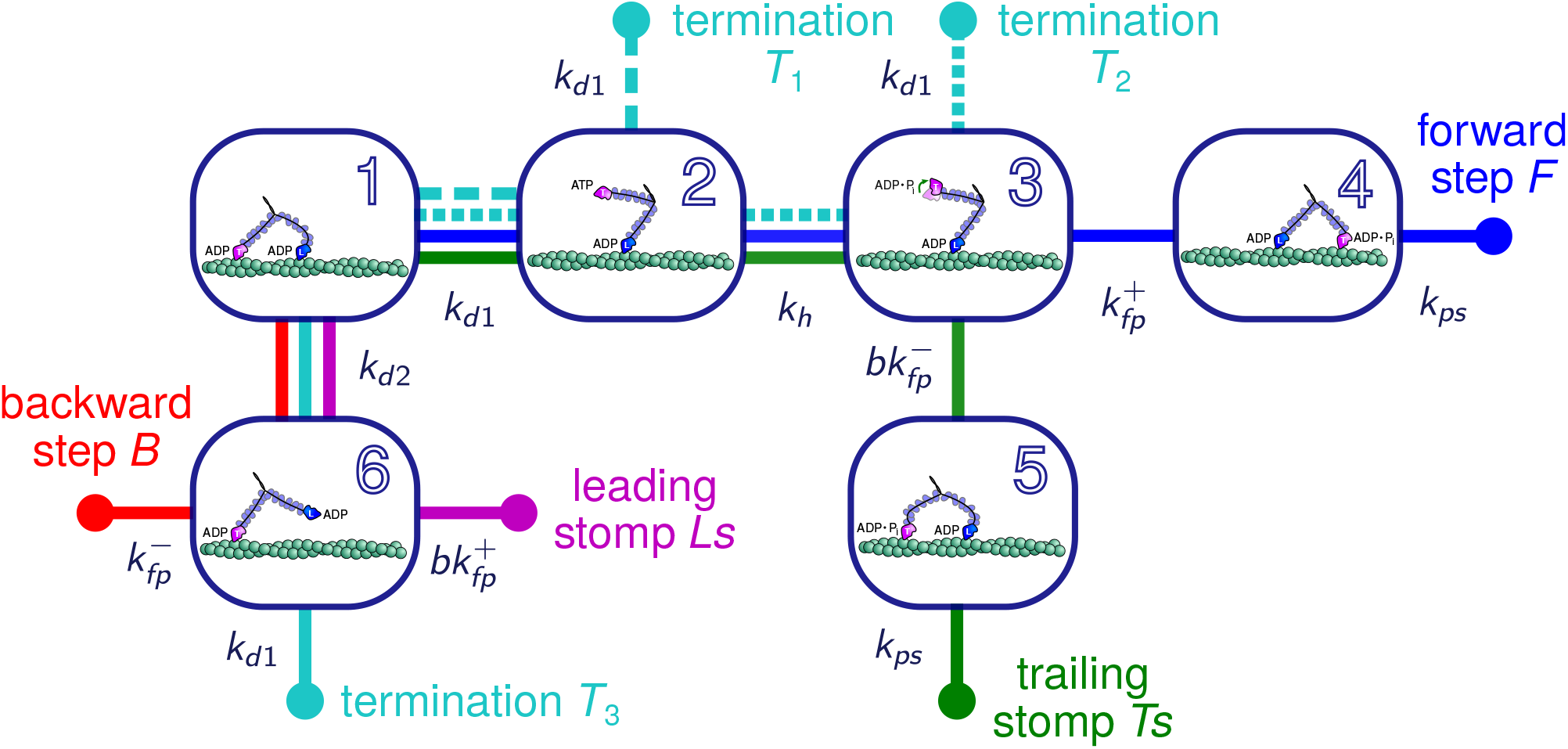
Kinetic network of myosin-V-like dynamics. Boxes 1-6 represent states, and lines connecting states represent pathways corresponding to the events listed in Tab. 1. Transitions between states are labeled by their rates *k*_*α*_. Myosin heads are marked T (trailing, pink) and L (leading, blue), denoting the initial positioning of the heads in state 1, where all the pathways originate.

Two pieces of information define each state of the kinetic network: the chemical state of the two myosin heads and which of the heads are bound to actin. The motor starts in the resting state (state 1), where both heads are attached to actin with ADP bound. To proceed, one head must detach. Notably, the detachment rates of the trailing and leading heads from the resting state (*k*_*d*1_ and *k*_*d*2_, respectively) are not the same since the leading leg experiences approximately 5 pN of backward strain^22^. This strain deforms the leading head to occlude the nucleotide channel, slowing ADP release. Thus, the leading leg detaches slower than the unstrained trailing head such that *k*_*d*1_ > *k*_*d*2_, which is responsible for a motor’s biased directionality^8^. In fact, ADP release is slowed so much that the dominant leading head detachment mechanism is ADP and ATP independent^8^.

There are several pathways that the motor can follow from the resting state, each defined by a sequence (*S*) of transitions between states (Tab. 1). The forward step (*S*_*F*_) and backward step (*S*_*B*_) begin with the trailing or leading head detaching from actin, respectively, and end with the motor moving ±1 step along the track. Our model assumes that the step size is of a constant length Δ. During the stepping events, the free head must diffusively find the forward (+) or backward (−) actin site with a mean rate of first passage 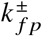. Since the leading head detaches without releasing ADP and remains in the post-powerstroke state, it can immediately bind to the actin site to complete the backward step. The trailing head, on the other hand, must first hydrolyze ATP with rate *k*_*h*_ and then release phosphate to initiate the powerstroke with rate *k*_*ps*_ once bound to actin. Doing so finishes the forward step.

**Table 1.**
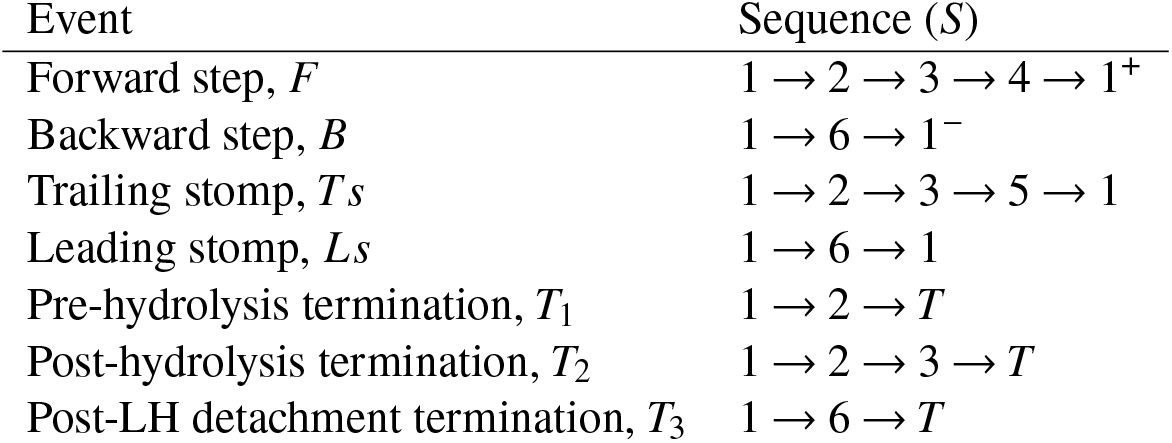
Events and their state sequences.

| Event | Sequence ($S$) |
| --- | --- |
| Forward step, $F$ | $1 \rightarrow 2 \rightarrow 3 \rightarrow 4 \rightarrow 1^+$ |
| Backward step, $B$ | $1 \rightarrow 6 \rightarrow 1^-$ |
| Trailing stomp, $T_s$ | $1 \rightarrow 2 \rightarrow 3 \rightarrow 5 \rightarrow 1$ |
| Leading stomp, $L_s$ | $1 \rightarrow 6 \rightarrow 1$ |
| Pre-hydrolysis termination, $T_1$ | $1 \rightarrow 2 \rightarrow T$ |
| Post-hydrolysis termination, $T_2$ | $1 \rightarrow 2 \rightarrow 3 \rightarrow T$ |
| Post-LH detachment termination, $T_3$ | $1 \rightarrow 6 \rightarrow T$ |

While in the diffusive search states, the free head could alternatively find the same actin-binding site that it came from. Such events result in either a trailing stomp (*S*_*Ts*_) or a leading stomp (*S*_*Ls*_) and no displacement along actin. Rebinding to the same site, however, incurs an additional energetic cost. The bound leg has to flex backwards in the case of the trailing stomp, while geometric hindrance inhibits the rebinding of the free leg in the post-powerstroke state for a leading stomp^8^. Our model incorporates these costs by using a binding penalty factor, 0 < *b* < 1 that modulates the first passage rates in the trailing and leading stomps, 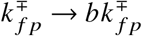, respectively.

A motor’s run then consists of any combination of stepping and stomping events until it ends with one of three terminating events 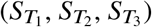. These occur when the bound head detaches before the free head binds to an actin site during a diffusive search. Since the bound leg is unstrained, all transitions to the terminated state from a diffusive search occur with the rate *k*_*d*1_.

### Polymer structure model of myosin

To calculate the first passage rates during the diffusive searches, we apply a previously developed polymer model of myosin-V (Fig. 2)^20^. As before, we model the neck domains of the motor as semi-flexible polymers with persistence length *l*_*p*_ and contour length *L*. If *l*_*p*_ ≫ *L*, the neck domains can be approximated as rigid rods. Our model assumes that the free leg rotates freely about the central junction during a diffusive search. At the bound head, the post-powerstroke state establishes a constraint angle *θ*_*c*_ between the attached neck and the axis of the actin track. The strength of the constraint angle is enforced by *v*_*c*_. In the limit *v*_*c*_ → ∞, the angle never deviates from *θ*_*c*_. Furthermore, the model incorporates the effects of an external resistive force applied to the motor’s junction with magnitude *f* and angle *θ*_*f*_ relative to the actin track.

**Figure 2.**
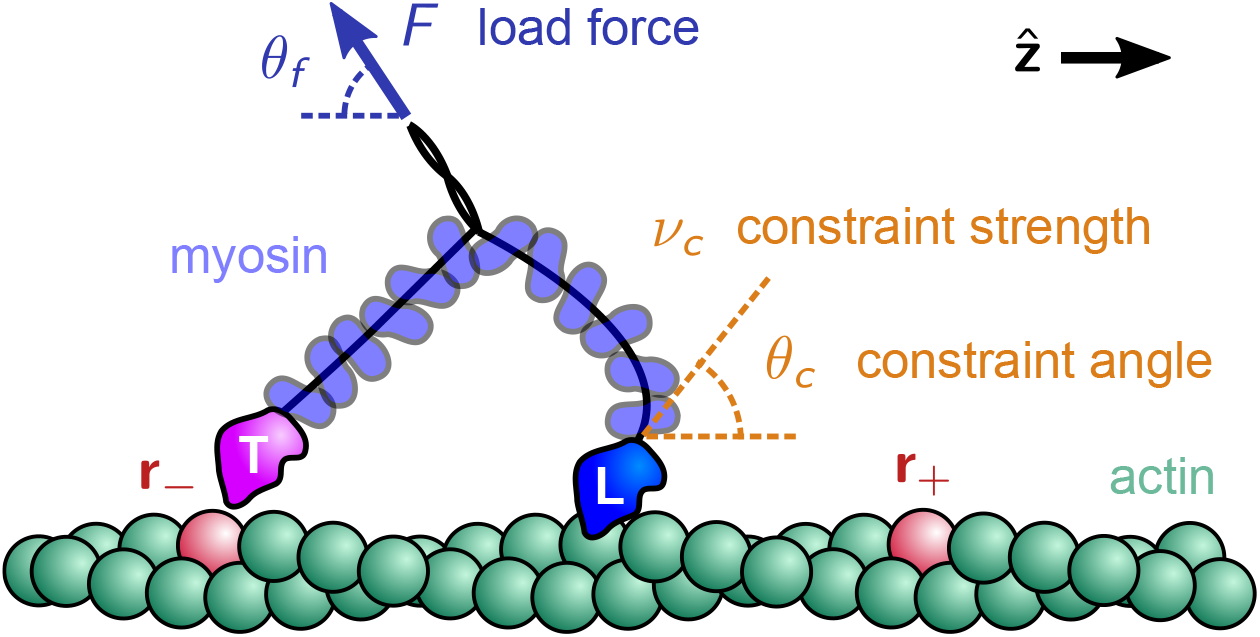
Schematic model of myosin-V-like motors. The actin filament is oriented along the 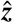 axis. The two myosin heads are marked T (trailing, pink) and L (leading, blue) based on their initial positions on the actin. The T head detaches from position ***r***_−_ and carries out a diffusive search for the forward binding site ***r***_+_. The dynamics of this search are influenced by two factors: i) the post-powerstroke leg constraint of strength *v*_*c*_, which favors a leading leg angle of *θ*_*c*_; ii) a load force of magnitude *F*, oriented at angle *θ*_*f*_ with respect to the 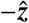 axis.

Previous work shows how these structural parameters determine the force-dependent diffusion space of the free head, *P*(**r)**, where **r** is the vector from the bound head to the free head^20^. This distribution can further be used along with the diffusivity of the myosin head, *D*_*h*_, and the capture radius, *a*, to calculate the mean rate of first passage to the forward or backward actin-binding sites located at **r**_±_:

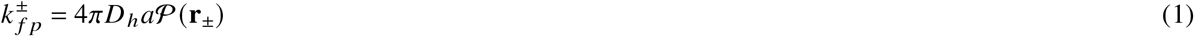

We report the intermediate expressions used to calculate Eq. 1 in the SI while a complete derivation may be found elsewhere^20, 21^.

We note that a recent extension of this polymer model addressed experimental evidence suggesting that the junction between the two legs instead undergoes constrained diffusion and included steps of varying size^21^. The updated model supported the constrained diffusion hypothesis, but the main observables relevant to the present work (for example velocities and run lengths) were essentially the same as in the simpler unconstrained model. We will thus continue to use the free diffusion model in this study.

### Event based picture of myosin dynamics

All seven events define a set, *E* = {*F, B,T s, Ls,T*_1_,*T*_2_,*T*_3_}, that entirely captures the motor’s dynamics such that the motor is guaranteed to execute one of them after leaving the resting state. Now, one can conceptualize a motor’s trajectory along actin as a sequence of randomly selected events, each with a probability ℙ_*i*_ such that ∑_*i*∈*E*_ ℙ_*i*_ = 1. Using the Markovian properties of the kinetic network, the event probabilities are just that of following the necessary sequence of states.

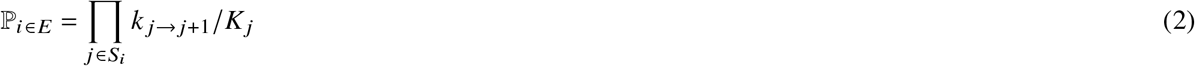

Where *k*_*j*→ *j*+1_ is the transition rate to the next state in the sequence and *K*_*j*_ is the total rate out of state *j*. Full expressions for the event probabilities in terms of the kinetic rates can be found in the SI. The set of events also offers an intuitive way to calculate the average number of events a motor completes during a run. As mentioned before, a run consists of any number of non-terminating events, followed by a single terminating event. Therefore, the number of events, *N*, follows a geometric distribution. Using 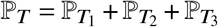, the probability of the motor completing *n* events is, ℙ_*N*_ (*n*) = (1 − ℙ_*T*_) ^*n*−1^ ℙ_*T*_ and has an average ⟨*N*⟩ = 1/ℙ_*T*_. In words, the average number of events is equal to the average number of times it takes for the motor to draw a terminating event.

One often wishes to narrow their focus to a subset of the motor’s events. A useful partition is between the stomping and non-stomping events. Doing so will help simplify the derivation of many subsequent expressions. Thus, we define the set of non-stomping events as Ê = {*F, B,T*_1_,*T*_2_,*T*_3_} such that its complement contains the stomping events, Ē = {*T s, Ls*}. If we restrict ourselves to only the events on one partition, the new event probabilities are simply the probabilities on *E* conditioned to the event belonging to the appropriate partition.

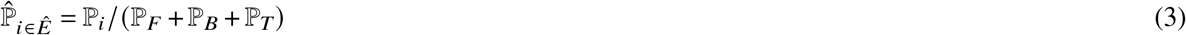

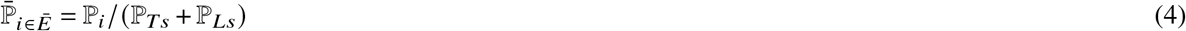

### Fitting to Myosin-XI

We now demonstrate that our generalized model captures the dynamics of myosins besides myosin-V. To do so, we fitted our model using a Bayesian approach to experimental distributions of dwell times and the detachment forces for single *Nt*. myosin-XI motors working against a compliant optical trap in the presence of 1 *µ*M and 100 *µ*M ATP^14^. We only used the dwell time distribution at 1 *µ*M ATP due to the limited temporal resolution of the optical trap system used in the original experiments as further explained in the SI^14^. To ensure the fitting procedure found motors with fast run velocities, a Gaussian prior using the experimentally determined mean run velocity and its standard error for myosin-XI in the optical trap at 100 *µ*M ATP (4.27 ± 0.35 *µ*m/s) was added. The Methods section contains additional details regarding the fitting procedure and data prepossessing steps.

We performed two versions of the model fit. Both used previously determined experimental values for myosin-XI’s *L*, Δ, and *k*_*h*_^14,23^. Values for myosin-XI’s *l*_*p*_, *D*_*h*_, *a*, and *b* in both fitting versions were assumed to be the same as myosin-V due to their high levels of structural homology^20^. In the first version of the fit, myosin-XI’s *θ*_*c*_ and *v*_*c*_ were also assumed to be the same as myosin-V’s so that only *k*_*d*1_ and *k*_*d*2_ were left as free parameters^20^. The second version of the fit allowed *θ*_*c*_ and *v*_*c*_ to vary as free parameters in addition to *k*_*d*1_ and *k*_*d*2_. Since *k*_*d*1_ is ATP dependent, we allow it to take different values at different ATP concentrations. By contrast, *k*_*d*2_ is ATP independent and was forced to have the same value at both ATP concentrations. Finally, we took *k*_*ps*_ as infinite since we assumed that the time it takes myosin-XI to complete the power stroke is much faster than any other kinetic step in the network. Tab. 2 reports the values of fixed and free model parameters for both sets of model fits. Fig. 3a-c compares the fitted and experimental distributions. The four parameter achieved a significantly better fit to the experimental data (Log posteriori of four parameter fit: -2422, two parameter fit: -2651). The model’s distribution of detachment forces at 1 *µ*M ATP varied the most between the two fits. Fitting with four free parameters better captured the experimental distribution’s main peak and tail Fig. 3b. Additionally, the four parameter fit more accurately replicated the experimentally reported average run velocity (⟨*V*⟩) of 4.27 *µ*m/s compared to the two parameter fit (⟨*V*⟩)four parameter fit: 4.37 *µ*m/s, two parameter fit: 6.80 *µ*m/s).

**Table 2.** Best fit model parameters for *Nt*. myosin-XI.

| Parameter | Two Param Fit | Four Param Fit | Notes |
| --- | --- | --- | --- |
| Leg contour length, $L$ (nm) | 33 | 33 | Ref. 14 |
| Leg persistence length, $l_p$ (nm) | 310 | 310 | Ref. 6 |
| Head diffusivity, $D_h$ (cm<sup>2</sup>s<sup>-1</sup>) | $5.7 \times 10^{-7}$ | $5.7 \times 10^{-7}$ | Refs. 24,25 |
| Constraint angle, $\theta_c$ (°) | 60 | 33 | Ref. 20, fit to experiment |
| Constraint strength, $\nu_c$ | 184 | 247 | Ref. 20, fit to experiment |
| Binding site separation, $\Delta$ (nm) | 35 | 35 | Ref. 14 |
| Capture radius, $a$ (nm) | 1 | 1 | Ref. 20 |
| Binding penalty, $b$ | 0.065 | 0.065 | Ref. 20 |
| Hydrolysis rate, $k_h$ (s<sup>-1</sup>) | 1000 | 1000 | Ref. 23 |
| TH detach rate (1 μM ATP), $k_{d1}^{1\mu M}$ (s<sup>-1</sup>) | 0.5 | 1.08 | Fit to experiment |
| TH detach rate (100 μM ATP), $k_{d1}^{100\mu M}$ (s<sup>-1</sup>) | 293 | 188 | Fit to experiment |
| LH detach rate, $k_{d2}$ (s<sup>-1</sup>) | 5.7 | 57 | Fit to experiment |

**Figure 3.**
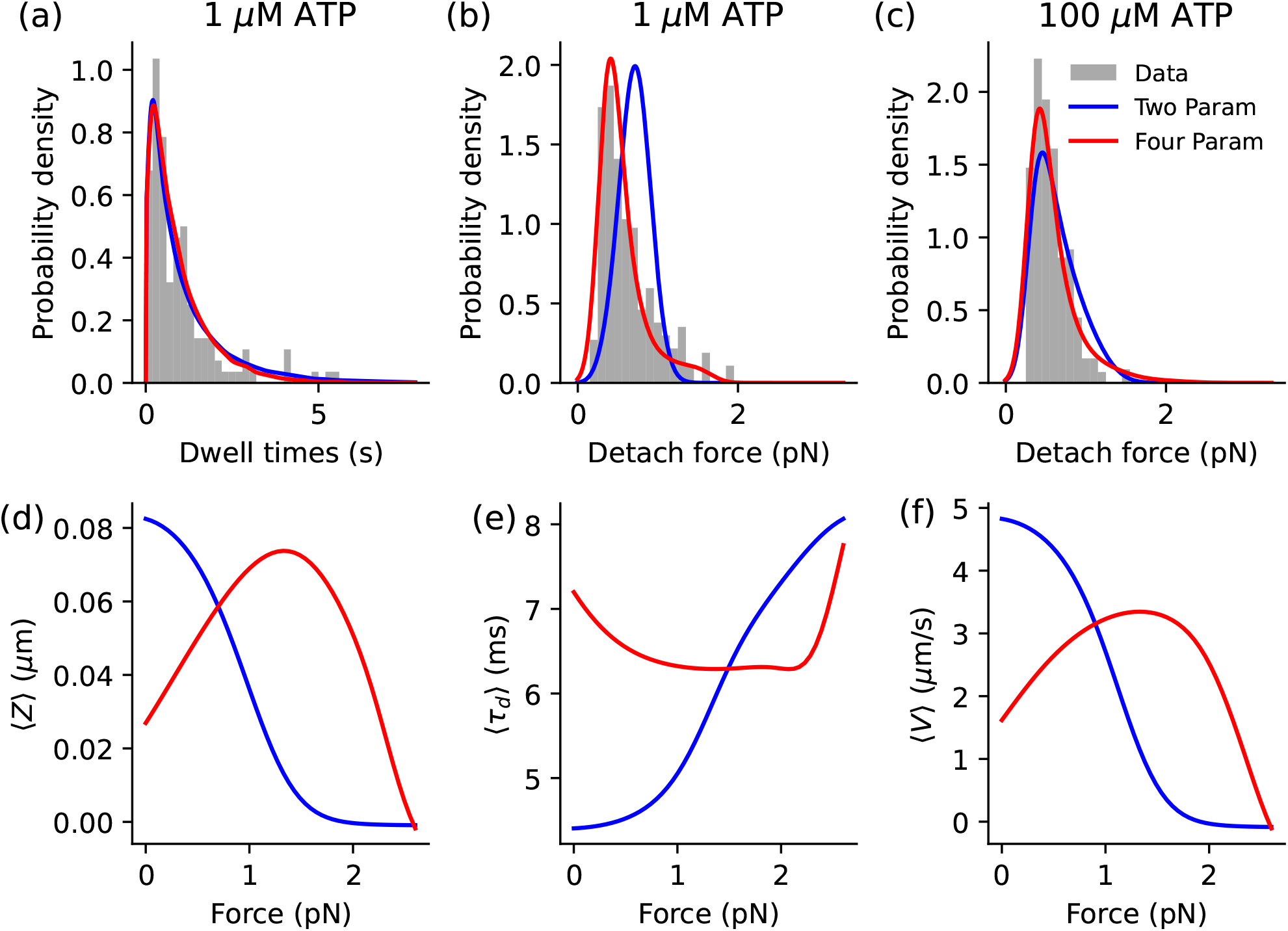
Fitting results versus experimental distributions and force dependent behavior. Three experimental distributions (gray) were used for fitting: (a) dwell times and (b) detachment forces at 1 *µ*M ATP, and (c) detachment forces at 100 *µ*M ATP for *Nt*. myosin-XI. Experimental data were obtained from ref^14^. The two parameter fit (blue) is the best fit when only *k*_*d*1_ and *k*_*d*2_ are allowed to vary as free parameters. The four parameter fit (red) is the best fit with *k*_*d*1_, *k*_*d*2_, *v*_*c*_, and *θ*_*c*_ as free parameters. The force dependence of the (d) average run distance, (e) average dwell time, and (f) average run velocity for motors according to the the two and four parameter fits while working against constant resistive load.

While the four parameter fit agreed more with the experimental data, further investigation revealed unexpected force-dependent behavior of the hypothetical motor of best fit (Fig. 3d-f). In particular, the average run distance (⟨*Z*⟩) and ⟨*V*⟩ first increase with resistive load before decreasing after 1.33 pN (Fig. 3d, f). One would typically expect ⟨*Z*⟩ and ⟨*V*⟩ to decrease with load. The average dwell time (⟨*τ*_*d*_⟩) also shows unexpected force-dependence where it first decreases before increasing after 2 pN (Fig. 3e). By contrast, the hypothetical motor of best fit from the two parameter fit demonstrates the typically expected force-dependent behavior for ⟨*Z*⟩, ⟨*V*⟩, and ⟨*τ*_*d*_⟩ where all observables show monotonic force dependence.

Such behavior is likely due to differences in *θ*_*c*_. The more acute *θ*_*c*_ found in during the four parameter fit (33^°^) versus that used for the two parameter fit (60^°^) places the diffusion space of the free head beyond the forward actin-binding site at low force. Larger forces on the motor pull the free head back to align its diffusion space with the actin-binding site. This increases the probability of completing a forward step and 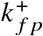 (Fig. S1) to increase ⟨*Z*⟩, decrease ⟨*τ*_*d*_⟩, both of which increase ⟨*V*⟩. Finally, large forces pull the free head away from the actin-binding, reducing 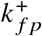 to decrease ⟨*Z*⟩, increase ⟨*τ*_*d*_⟩, and slow ⟨*V*⟩. While such a motor is theoretically possible, additional experiments are needed to determine whether myosin-XI behaves more like to motor predicted by the four parameter fit or the two parameter fit.

Regardless, we have shown that our model can reasonably explain the dynamics of motors besides myosin-V. Thus, we go on to derive general expressions for observable quantities describing the dynamics of any myosin-V-like motor under a constant resistive load and interrogate the nature of their overarching design space.

### Run distances

A critical experimental quantity that characterizes a molecular motor is the net distance it moves during a run. Here we define the run distance under a constant resistive load, *Z*, as the number of steps the motor displaced from its starting position on actin at the time of termination.

The event-based description of myosin dynamics offers a straightforward way to calculate the average run distance; multiply the average displacement over the non-stomping events in *Ê* by the average number of non-stomping events, 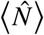. Like the number of events in general, 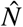 is geometrically distributed but with 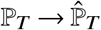 and average 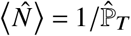. Since forward and backward stepping events move the motor ±1 step, respectively, and the terminating events do not change the motor’s position, the average run distance is,

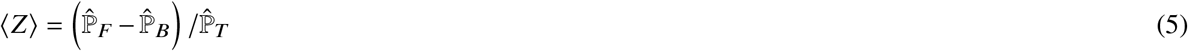

Note that the normalizing factor used to calculate the 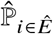 (Eq. 4) cancels in the numerator and denominator of Eq. 5. Therefore, the average run distance is the same if one instead uses the probabilities on *E*.

Next, we derive an expression for the distribution of run distances, ℙ_*Z*_ (*z*). Since our model allows for back stepping, an infinite number of trajectories can result in a single run distance. Thus, the probability that a motor terminates at a certain distance from its starting location is the sum of all trajectories producing the desired displacement. In terms of the simplified set of events,

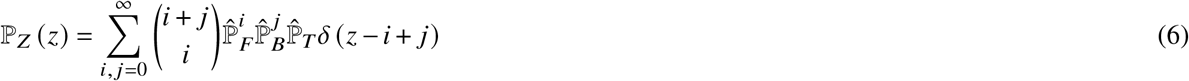

where the first factor in parentheses is the binomial coefficient, and *δ* is the Kronecker delta function. As shown in the SI, evaluating the double sum for the case *z* ≥ 0 results in an expression in terms of the Gaussian hypergeometric function,_2_*F*_1_ (*a, b*; *c*; *x*), with parameters *a, b, c* and argument *x*,

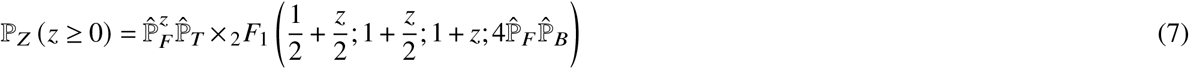

We further explain in the SI that interchanging *F* and *B* with each other handles the case for *z* < 0. The hypergeometric function is of the form_2_*F*_1_ (*a*; 1/2 + *a*; 2*a*; *x*) and has a closed form solution (Eq. 15.1.14 of ref 26),

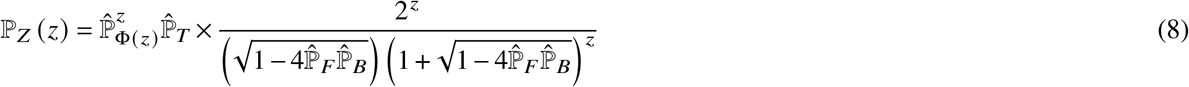

where Φ(*z* ≥ 0) = *F*, Φ(*z* < 0) = *B* to indicate directionality.

In this form, the 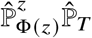 factor is the same as the probability that the motor runs *z* consecutive steps in the direction of z (forwards if positive, backwards if negative) and then detaches, and represents the minimum number of displacing events needed to achieve a run distance of *z*. Thus, the hypergeometric function encodes information from all the other trajectories that could have resulted in the same run distance and scales the probability of the most efficient trajectory accordingly. Also, note that our result for the distribution of run distances is a generalization of a similar expression previously found for a simpler periodic one-state model of motor dynamics^27^.

### Average dwell times and run times

An equally important quantity is how long a motor spends at one position on actin before moving or terminating. In our model, we define the motor’s dwell time, *τ*_*d*_, as the amount of time it spends stomping in place plus the time it takes to complete the single non-stomping event needed to move between actin sites. Mathematically, we can express the dwell time in terms of the time it takes to complete individual events, *τ*_*i∈E*_.

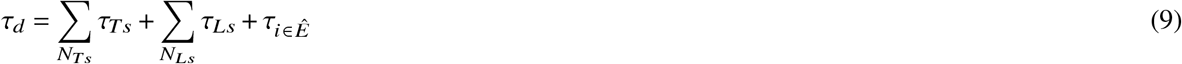

*N*_*Ts*_ and *N*_*Ls*_ are the random numbers of trailing and leading stomps that happen before the transition event.

As detailed in the SI, the average dwell time can then be expressed in terms of the average event-times, ⟨*τ*_*i* ∈*E*_⟩, the event probabilities on the set partitions *Ê* and *Ē*, and the average number of total stomps before the transition event, 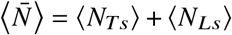.

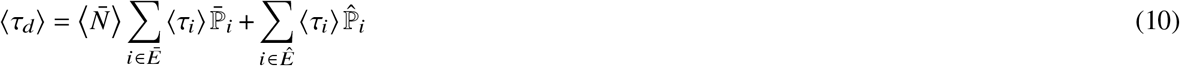

In this form, the first term accounts for the average time spent stomping, and the second term is the average time spent transitioning. One can further express 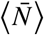 in terms of event probabilities by considering how the transition event ends the preceding sequence of stomps. Thus, 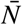 is geometrically distributed with average,

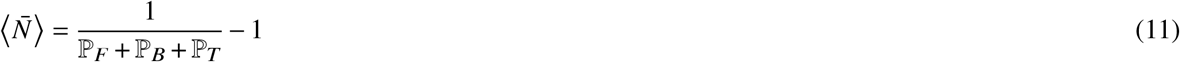

Note that these probabilities are on the complete set of events, *E*.

To calculate the average event times, recall that each event is simply a sequence of states with exponential wait times. Thus, the event times formally follow a hypoexponential distribution with an average equal to the sum of the average wait times of each state.

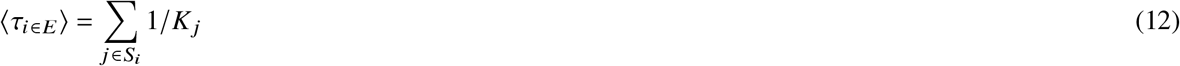

As in Eq. 2, *K*_*j*_ is the total transition rate out of state *j*. The SI reports full expressions for all the average event times in terms of the kinetic rates.

With the average dwell time in hand, an expression for the average run time quickly follows. First, we define the run time, *τ*_*r*_, as how long a motor spends with at least one leg attached to actin. Then, since a run consists of 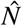 transition events, the run time e qua ls to the sum of 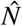 dwell times, 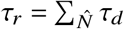. The average run time is then just the average number of transitions, 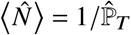 multiplied by the average dwell time,

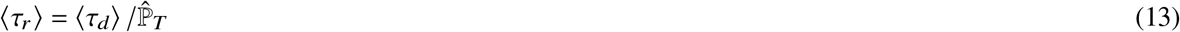

The average run time can alternatively be expressed as the sum of *N* event times on the complete set *E*, which we report in the SI.

### Distribution of dwell times and run times

However, finding insightful mathematical expressions for the distributions of dwell and run times poses more of a challenge. Both the dwell times and run times follow phase-type distributions. A phase-type distribution characterizes the arrival times, *τ*, to a single absorbing state on an arbitrary Markovian network. Two pieces of information parameterize a phase-type distribution. One is the network’s transition matrix, Ω. We use the convention where the off-diagonal elements of the transition matrix Ω_*i, j*_,*i* ≠ *j* are the transition rates, *k*_*j*→*i*_. The diagonal elements enforce the property that the columns of the transition matrix must sum to zero: Ω_*i,i*_ = − ∑_*j*≠*i*_ *k*_*i*→ *j*_. In the case of our kinetic network, the transition matrix for the dwell times, Ω_*d*_, has dimensions of 7 × 7 where the last state of each event-sequence of *S*_*i ∈ Ê*_ has been replaced with an absorbing state. Furthermore, the absorbing state occupies the last column of the transition matrix so that Ω_*d*_ can be partitioned into a 6 × 6 submatrix, *A*_*d*_, a column vector of zeros, **0**, and a row vector, **a**_**d**_ = −**1***A*_*d*_ where **1** is a vector of ones.

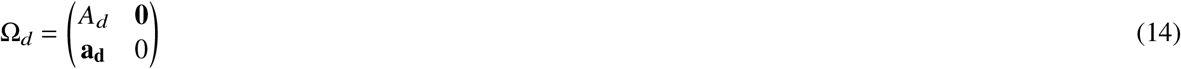

The second parameter needed to define a phase-type distribution is a probability vector, **p**_0_, representing the chance of a random walker staring in any of the network’s states. Since the motor always starts in the resting state (state 1), the initial probability vector has six elements with a value of one at the first index and the rest as zeros. The SI reports the full expressions of Ω_*d*_ (Eq. S51) and **p**_0_ (Eq. S52).

The density function for the distribution of dwell times can then be derived from the general solution to the master equation that dictates how the probability of being in a given state on the network evolves in time^28^.

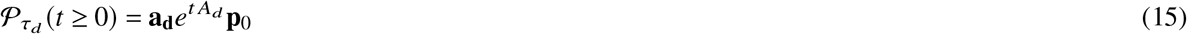

where 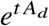 the matrix exponential. When *t* < 0, the distribution becomes zero. For notational simplicity, we will assume that the domain for all distributions involving time is implied. One can also express the average dwell time and its variance, 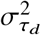 in terms of **p**_0_ and the matrix inverse of *A*.

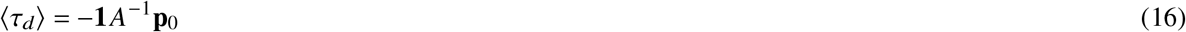

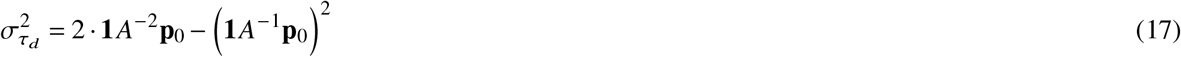

Note that ⟨*τ*_*d*_⟩ calculated here is equivalent to that used in Eq. 10 but packaged with matrix notion.

One approach to get the run time distribution is to define a new transition matrix, Ω_*r*_, where the only difference from Ω_*d*_ is that only the terminating events connect to the absorbing state. In other words, the motor gathers time stepping and stomping around the network until both legs detach from actin. Then, one simply gets another distribution of the same form as Eq. 15. We report the necessary transition matrix in the SI (Eq. S53).

### Gamma approximation of the dwell time distribution

While the phase-type representations of the dwell time and run time distributions are exact and extremely compact, they will not be suitable for our upcoming analytical purposes. In particular, we will want to represent the run time distribution in terms of a sums of dwell times when deriving expressions relating to a motor’s run velocity and take the limit of those sums to infinite terms. As we detail in the SI, a phase-type distribution representing a infinite sum of dwell times has a infinite dimensional transition matrix. Thus, we aim to approximate the dwell-time’s phase-type distribution using a more well behaved distribution.

Of the many candidate distributions, the gamma distribution best serves our needs. We will use the convention where the gamma distribution is parameterized with shape parameter *α* and scale parameter *θ*, and has a density function,

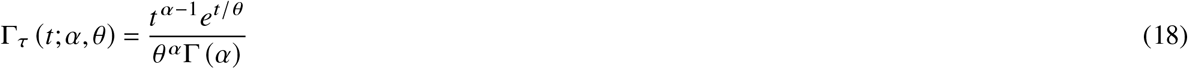

where 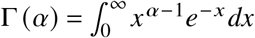 is the gamma function. To calculate the approximating shape and scale parameters, we match the gamma distribution’s mean, *αθ*, and variance, *αθ*^2^, to those of the dwell time’s phase-type distribution, then solve for *α* and *θ*.

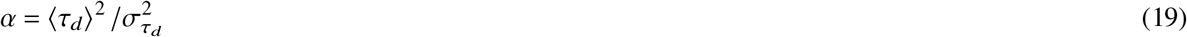

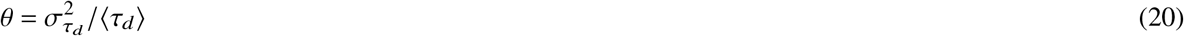

Using the gamma distribution and moment-matching method comes with many important advantages. First and foremost is the valuable property that the sum of 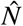 dwell times approximated by the gamma distribution is another gamma distribution with shape parameter 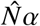 and the same scale parameter. Now, instead of needing infinite matrices to represent infinite sums of dwell times, we only need to take the limit of a scalar. Furthermore, average values that depend on the dwell time remain exact because the moment-matching method guarantees that the approximation shares the original mean.

Finally, the gamma distribution reduces to an Erlang distribution when *α* is a positive integer, representing the arrival times from a sequence of *α* identical states with *θ* average waiting times. Since the Erlang distribution is a subclass of the phase-type distribution, the gamma approximation essentially recasts the original network into the simpler sequence of states with the added flexibility of non-integer *α*. While we do not formally prove that gamma approximation becomes exact for positive integer values of *α*, numerical comparisons between the actual dwell time distribution and the gamma approximation for a random sampling of myosin-V-like motors show that it is excellent in these cases (Fig. 4). Additionally, the calculated *α* for this sampling reveal that most approximating shape parameters are close to either one or two (Fig. 4b). Finally the motor from the sampling where the approximation performed the worst still showed good agreement with the exact phase-type distribution(Fig. 4c). Subsequent results will show how this property of *α* offers novel insight into the design space of motors. Together, these establish the efficacy of the gamma approximation for the phase-type distribution of dwell times.

**Figure 4.**
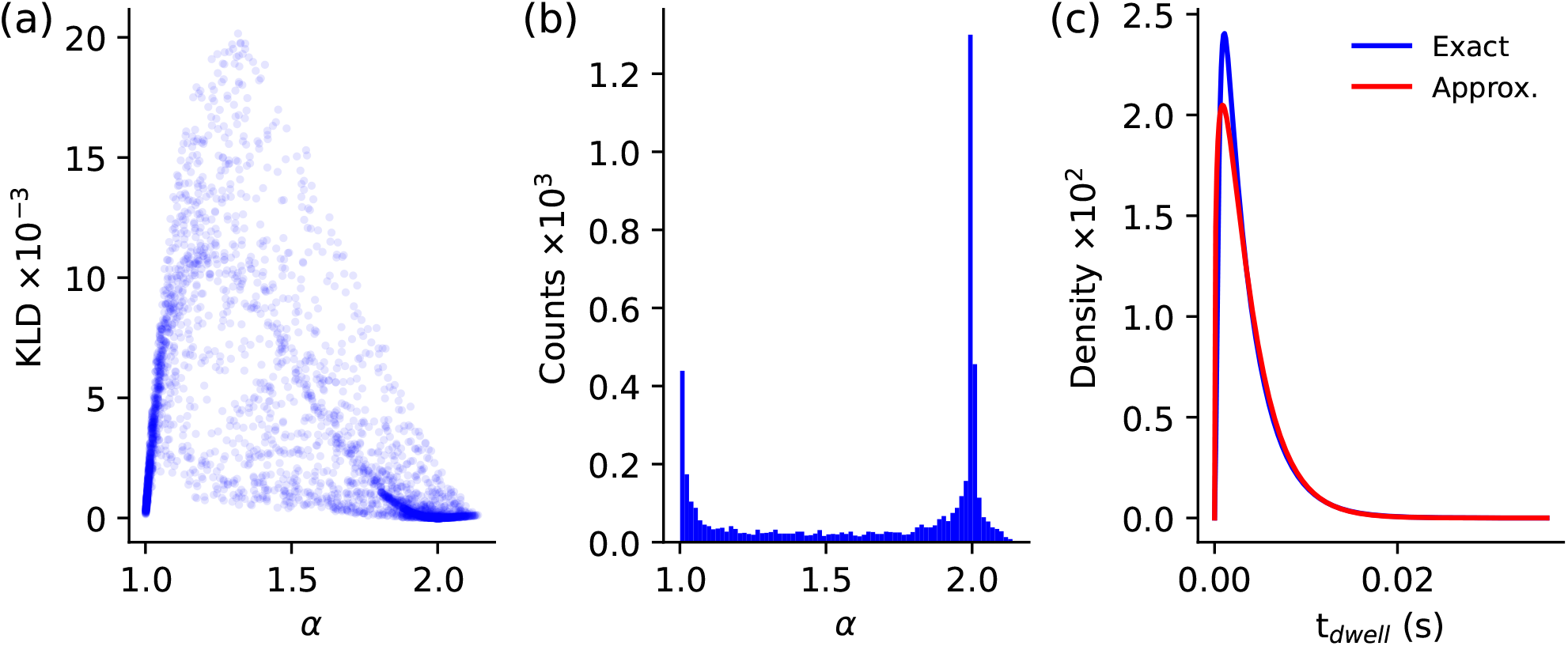
Comparison between the dwell-time phase-type distribution and its gamma distribution approximation. (a) Scatter plot of the Kullback-Leibler divergence (KLD) between the true and approximate dwell-time distributions as a function of the approximating shape parameter (*α*) for 5000 random motors. The values for *k*_*d*1_, *k*_*d*2_, *k*_*h*_, 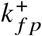, and 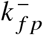 were randomly drawn between 10 and 10^6^on a log-uniform distribution. Motors with *α* = 1 and *α* = 2 have near perfect agreement (KLD = 0) between approximate and exact distributions. (b) Histogram of *α* parameters showing clustering around *α* = 1 and *α* = 2. (c) Comparison between the exact (blue) and approximate (red) dwell time distributions for the motor in the sampling with the highest KLD.

### Run Velocities

We now have all the necessary tools to derive quantities describing a myosin-V-like motor’s run velocity, defined as the run distance divided by the run time: *V* = *Z* /*τ*_*r*_.

We start by deriving the distribution of run velocities. The most general form of the velocity distribution is defined in terms of the joint density of run times and run distances, 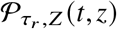. Here, 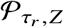 is a mixed density since the run time is a continuous random variable while the run distance is discrete. To get the distribution’s value at a specific run velocity, *v*, sum all parts of the joint density such that *v* = *z*/*t*,

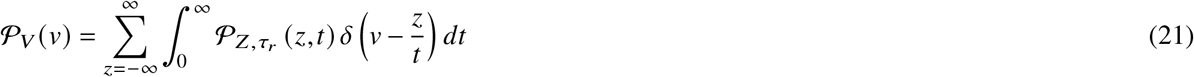

where *δ* is the Dirac delta function. For *v* ≠ 0, one can evaluate the time integral with a change of variables to get a single sum over run distances. When *v* = 0, the distribution has a delta contribution with mass equal to the probability that the run distance is zero. By Eq. 8, ℙ_*Z*_ (0) = ℙ_*T*_ (note that there is no hat) and we obtain

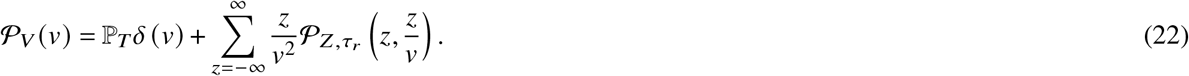

Getting the run velocity distribution in terms of known quantities requires a functional form for the joint distribution of run times and run distances. We construct an appropriate expression by considering that trajectories consisting of *i* forward steps, *j* backward steps, and a single terminating event have a total of *i* + *j* + 1 transition events along actin. Thus, the run times for these trajectories, *τ*_*i*+ *j*+1_ is the sum of *i* + *j* + 1 dwell times. Using the gamma approximation for the dwell time, the distribution of *τ*_*i*+ *j*+1_ is 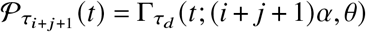. Noting that *Z* = *i* − *j* and using the probability of observing such a trajectory, we can write the joint distribution as

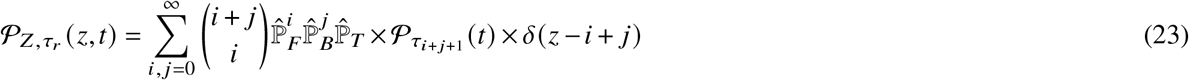

where *δ* is the Kronecker delta function. Substituting Eq. 23 into Eq. 22 completes the derivation of the run velocity distribution.

In the SI, we further calculate the average of run velocity, ⟨*V*⟩ = ⟨*Z*⟩/⟨*τ*_*r*_⟩, in terms of a generalized hypergeometric function,

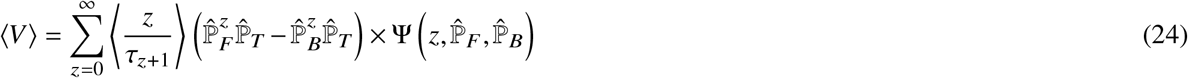

where ⟨*z*/*τ*_*z*+1_⟩ = *z*/[(*z* + 1) *αθ* − *θ*] is the fir st negati ve moment of the run time for a trajectory consisting of *z* + 1 transitions multiplied by *z*, and Ψ 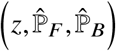 is a generalized hypergeometic function of the form, 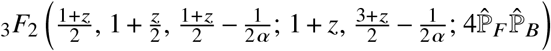.

As in the expression for distribution of run distances, runs consisting of purely forwards or purely backward steps play a central role. For each term in the sum over *z*, the first factor calculates the average run velocity for a unidirectional run that displaces the motor *z* steps in either direction. The second factor weights this velocity according to the probability of the forward and reverse trajectories. Analogous to Eq. 8, the hypergeometric function encodes the contributions from all the other trajectories that result in the same run velocity by running farther in less time or vice versa. Furthermore, unlike the density function, this expression for the mean is exact since it only depends on the average dwell time quantities.

Eq. 24 has a closed form when *α* = 1. We show in the SI that, in this case, the average run velocity reduces to the quotient of the average run distance and run time.

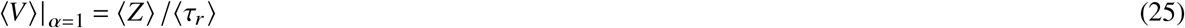

We were unable to simplify Eq. 24 for the *α* = 2 case, but it can be evaluated numerically.

Finally, it is important to note that the values calculated by Eq. 24 include the contributions from trajectories of zero run distance. Since in vitro measurements of motor dynamics cannot count these non-productive trajectories, average run velocities determined experimentally may significantly disagree with Eq. 24 in the cases where P_*T*_ is large. Fortunately, it is not difficult to determine the average run velocity conditioned on productive runs.

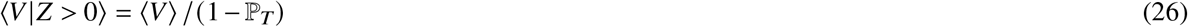

### The myosin speed-limit

With these velocity expressions, we now ask whether an intrinsic limit exists for how fast a myosin-V-like motor can run.

Considering the fastest motor, we can immediately apply some intuitive simplifications to the existing model. First, the fastest motor will have perfect gating (i.e. *k*_*d*2_ = 0). If it did not, the motor could take backward steps that add to the run time but subtract the run distance. Both reduce the run velocity. Setting *k*_*d*2_ = 0 also eliminates the possibility of leading stomps and the *T*_3_ termination event. These also decrease the run velocity by increasing run time without increasing run distance. Thus, perfect gating only allows the motor to make forward steps, trailing stomps, and terminate via the two remaining termination events. As detailed in the SI, we can find new expressions for the distribution of run velocities for motors with perfect gating, *V* ^′^, along with its average. We use primed quantities to denote signify perfect gating.

A second simplification we can make for the fastest motor is taking *k*_*ps*_ → ∞ since *k*_*ps*_ only increases the run time in our model. Additionally, we assume that the motor works against a constant force such that is fixed. 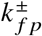 Now, *k*_*d*1_ and *k*_*h*_ are the only two free parameters left, which allows us to numerically determine the *k*_*d*1_ that maximizes ⟨*V* ^′^⟩ at a given *k*_*h*_ 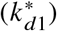. Calculating *α* when 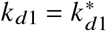, reveals that they all are roughly equal to two (Fig. S2). As shown in the SI, the new average run velocity takes a simple form in this case.

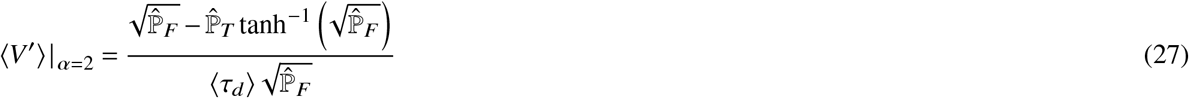

We also note that plotting 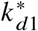 versus *k*_*h*_ on log-log scales suggests that the two share a power-law relationship, 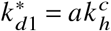 (Fig. 5a). The exponent is close enough to one half that we approximate the relationship with a square-root: 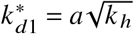. We found the same square root relationship for a wide range of motors suggesting it generally holds within the myosin-V-like design space (Fig. 5). Replacing *k*_*d*1_ in the underlying expressions for the probabilities and average dwell time in Eq. 27 with the square-root relationship leaves *k*_*h*_ as the final free parameter. Finally, taking the limit of *k*_*h*_ → ∞ results in the theoretical maximum average run velocity, ⟨*V*⟩^*\**^, that a myosin-V-like motor can achieve:

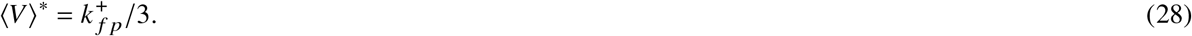

**Figure 5.**
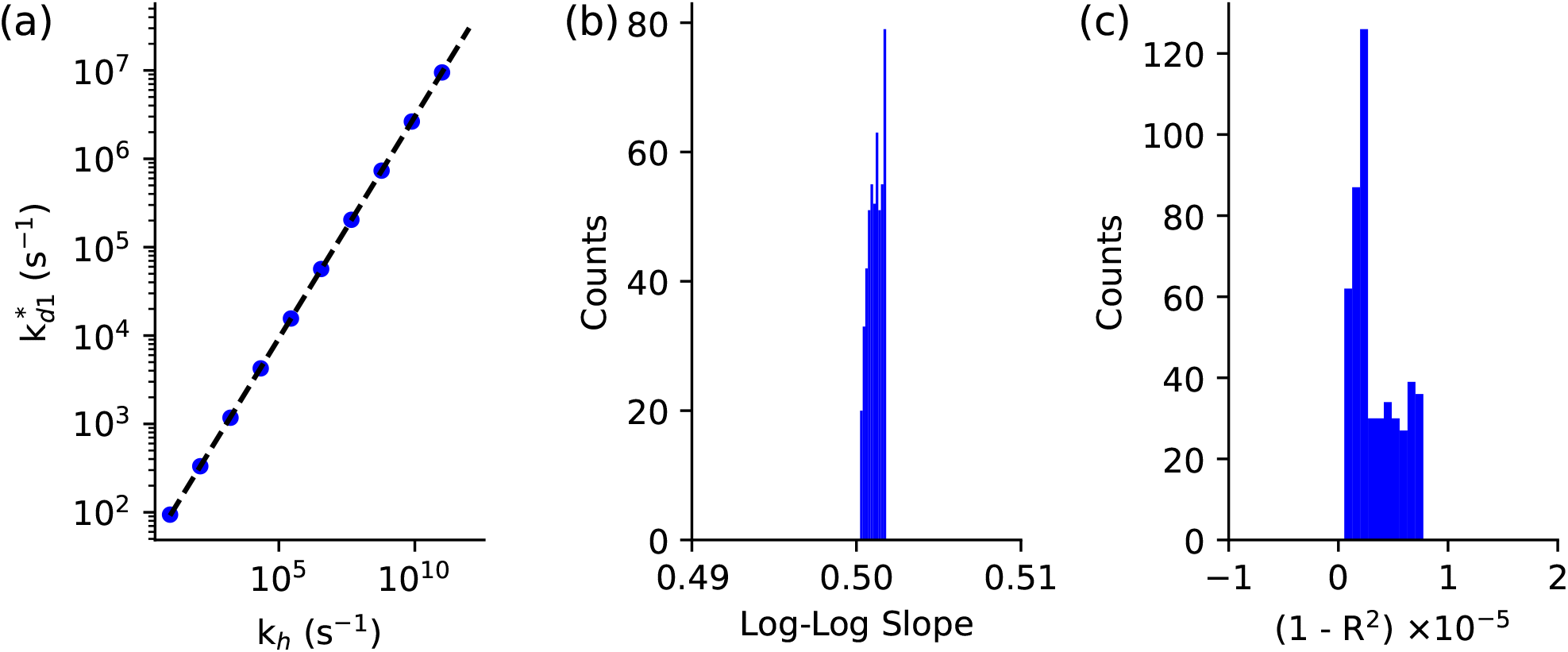
Empirical square root relationship for the k_d1_ that maximizes run velocity at a given k_h_. (a) Example log-linear fit (black dashed line) of *k*_*d*1_ values that maximize run velocity at a given *k*_*h*_ (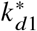, blue dots) for a motor with perfect gating and myosin-V structural parameters according to ref 20. (b) Histogram of the slopes of log-linear fits as in panel (a) for 500 motors with perfect gating and random 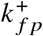 and 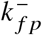 values between 10 and 10^6^ s^−1^ drawn from a log-uniform distribution. Results show that all slopes are approximately 0.5 (square root). (c) Histogram of 1 − *R*^2^ values for the log-linear fits in (b) showing the extremely high linearity of the relationship.

To confirm this limit, we plot the average run velocity for a wide variety of motor parameterizations versus 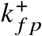 and show that all points remain below the line 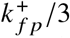 (Fig. 6a). We also color-coded the points by their approximating shape parameter to show that all the points very close to the speed limit have *α* ≈ 2. Note that being a motor with *α* ≈ 2 does not in itself guarantee that it will be at the speed limit, since several *α* ≈ 2 motors are far from the limit while some *α* ≈ 1 motors approach it.

**Figure 6.**
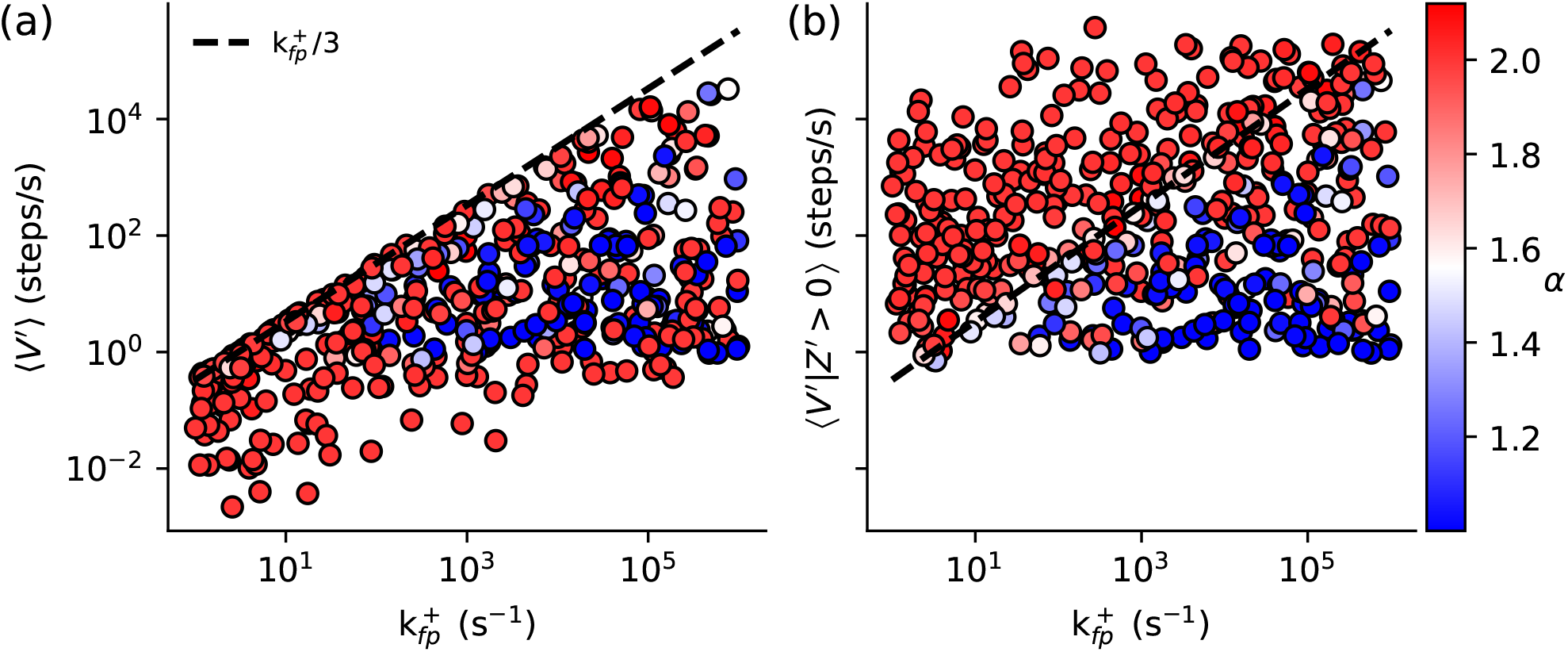
The absolute average speed-limit of myosin-V-like motors. (a) The average run velocity (⟨*V*^′^⟩) for 500 hypothetical myosin-V-like motors with perfect gating and random 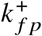 values. All velocities fall bellow the line 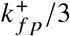 (black dashed line) as predicted. (b) The average run velocities in (a) when conditioned on productive runs (⟨*Z*^′^ > 0⟩). Motors are color coded according to their approximating shape parameter (*α*). Average velocities were determined from fifty thousand trials. The *k*_*d*1_, *k*_*h*_, 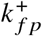, and 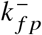 parameters of these motors were randomly selected from a log-uniform distribution between 10 and 10^6^ s^−1^.

Recall, however, that validating ⟨*V*⟩^*\**^ experimentally could prove difficult (assuming such a motor even exists) since it includes contributions from non-productive runs. Using Eq. 26 to calculate the average run velocity from productive runs only shows that ⟨*V* ^′^|*Z*^′^ > 0⟩ is likely unbounded (Fig. 6b). Mathematical, this is because P_*T*_ approaches one as 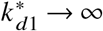 and makes the denominator of Eq. 26 infinitesimally small. Interestingly, however, only motors with *α* ≈ 2 break the original speed limit (Fig. 6b). This is likely because *α* ≈ 2 motors have a larger P_*T*_ than *α* ≈ 1 motors.

### The myosin Pareto frontier

While the maximum average run velocity offers an ultimate limit on how fast a motor can run, there may yet be interesting relationships between the velocity and other observable quantities at sub-maximal speeds. The apparent trade-off between run distance and run velocity seen experimentally is a prime candidate to investigate^13^.

Thus, we generated a scatter plot on log-log scales of average run velocity versus average run distance for a broad sampling of parameter choices and color-coded the points according to their approximating shape parameter (Fig. 7). We only investigated motors with perfect gating for two reasons. First, we are primarily interested in motors that maximize the relationship between average run distance and velocity. Back stepping would decrease both. Second, accurately calculating the average run velocity in the presence of back stepping becomes numerically unstable for motors with average run distances of over 700 steps (Fig. S3).

**Figure 7.**
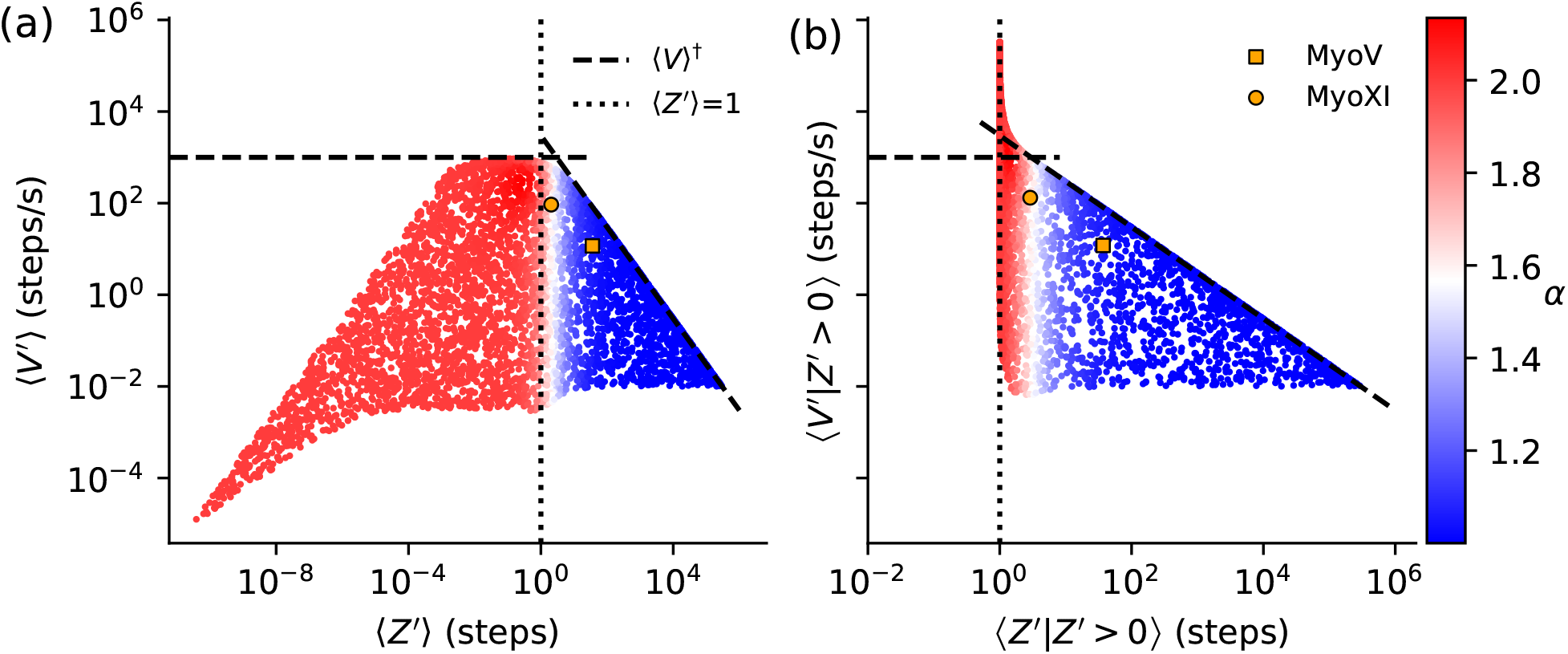
The average run velocity-distance Pareto frontier. (a) Scatter plot of average run velocities (⟨*V* ^′^⟩) and run distances (⟨*Z*^′^⟩) for 5000 motors with perfect gating, structural parameters like myosin-V^20^, and random values of *k*_*d*1_ and *k*_*h*_ drawn between 10^−2^ and 10^6^ on a log-uniform distribution. Points are color coded based on the value of their approximating parameter (*α*) and bounded by the Pareto frontier (⟨*V* ^†^⟩). The vertical dotted line at ⟨*Z*^′^⟩ = 1 separates the *α* = 1 (right, blue) and *α* = 2 (left, red) regions. The locations of myosin-V (MyoV^20^) and *Nt*. myosin-XI (MyoXI, four parameter fitting results at 100 *µ*M ATP) are shown as an orange square and circle, respectively. (b) The same data as in panel (a) but with the average run velocities and run distances conditioned on productive runs (*Z*^′^ > 0).

The results reveal two main features. First, a bounding curve limits the maximum average run velocity that a motor with a given average run distance can obtain (Fig. 7a). An initial glance suggests that the curve has three distinct regions: a central flat region flanked by two log-linear regions. In the SI, we find approximations for the bounding curve, ⟨*V*⟩^†^, in these regions and show that only the central and right regions are legitimate bounds:

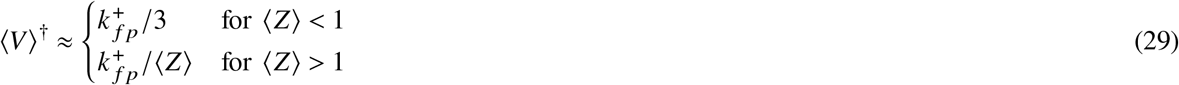

The log-linearity of the left region’s boundary is an artifact of the finite parameter range used to evaluate the model and, in reality, belongs to the central region. Fig. S3a shows that Eq. 29 holds when back stepping is allowed.

We also investigated the relationship between average run velocity and run distance when conditioned on productive runs (Fig. 7b). We show in the SI that the right side of the Pareto frontier is the same as in the unconditioned case. We also show in the SI that the asymptote on the left side occurs at ⟨*Z*^′^|*Z*^′^⟩ > 0 = 1. However, motors with non-perfect gating can break though to the left of the asymptote (Fig. S3b).

Eq. 29 defines a Pareto-frontier that dictates the best average run velocity that a motor with a given run distance can achieve. For the region where ⟨*Z*⟩ < 1, Eq. 29 approximately equals the theoretical speed-limit found in Eq. 28. Once ⟨*Z*⟩ becomes greater than one, the maximum average velocity obtainable falls proportionally with ⟨*Z*⟩. Therefore, the myosin-V-like motor that simultaneously maximizes its average run velocity and distance would exist near the division between the two domains.

The second observation from Fig. 7 is the stark division of approximating shape parameter values between the two regions. Motors with *α* ≈ 2 exclusively reside in the region where ⟨*Z*⟩ < 1 while *α* ≈ 1 motors lay on the other side with only a small transition zone between them. Seeing the approximating shape parameters in this light offers a key insight into their physical meaning. In essence, motors with *α* ≈ 2 are drag racers that can run fast but only for short distances, while motors with *α* ≈ 1 are long haulers that trade speed for endurance.

The equation for the average run distance reveals the mathematical reasons for why. When ⟨*Z*⟩ > 1, Eq. 5 implies that the effective probability of taking a forward step is relatively larger than the probability of terminating. Looking at the network in Fig. 1 and the equations for P_*F*_ (Eq. S8) and P_*T*_ (Eqs. S12, S13, S14) reveals that the rate *k*_*d*1_ must be smaller than both *k*_*h*_ and 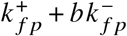. If it were not, then the motor would begin to favor the termination pathways over forward stepping. Thus, assuming a fast power-stoke and *k*_*d*1_ > *k*_*d*2_, *k*_*d*1_ and *k*_*d*2_ are the slowest rates such that the resting state has the longest escape time of the entire network. In the cases where a single state dominates the network, the dwell time distribution approximately becomes an exponential distribution equivalent to the gamma distribution with *α* = 1.

When *α* ≈ 2, Fig. 7 and Eq. 29 suggest that the motor is likely to terminate before taking a single step. Since there are multiple ways for the motor to terminate, multiple scenarios can result in *α* ≈ 2. First consider the case when *k*_*h*_ ≫ *k*_*d*1_. Here, the 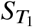 termination pathway dominates the network. In this limit, both states in 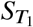 will have approximately the same rate *k*_*d*1_, and the dwell time distribution behaves like an Erlang distribution with a shape parameter of two (i.e., *α* = 2). Alternatively consider the scenario where 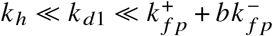. Now, the 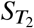 termination pathway dominates. Typically, the motor has to visit three states along 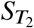. However, with *k*_*h*_ being much faster than *k*_*d*1_, the pre-hydrolysis state becomes insignificant, so only the resting and post-hydrolysis states contribute to the dwell time. As before, *k*_*d*1_ accounts for most of the total rate from both states meaning that the dwell time distribution again behaves like the Erlang distribution with two states.

## Discussion

By generalizing a theoretical framework developed to describe myosin-V dynamics^20, 21^, we have uncovered the theoretical design-space of all myosin-V-like molecular motor proteins. As a display of the generalized model’s efficacy, it predicted the dynamic behavior of the fast *Nt*. myosin-XI motor protein functioning in a compliant optical trap (Fig. 3). This is the first time a single model with a complete microscopic description of motor dynamics has explained experimental data from distinct myosin classes. Our fitting results revealed that the *Nt*. myosin-XI motor domain may form a stronger and more acute constraint angle with actin filaments compared to myosin-V (Tab. 2). Further investigation showed that these potential differences in actin interactions lead to some unintuitive force dependent behavior that allow *Nt*. myosin-XI to maximize run velocity and run distance while under a non-zero load (Figs. 3 and S1). If experiments validate our predictions, *Nt*. myosin-XI would be an intriguing example of a motor tuned to function within a specific force regime. Other fast members of the myosin-XI family may use a similar strategy.

In many respects, our theoretical results parallel general frameworks for motor dynamics recently developed for non-myosin-V-like motors, for example myosin-VI^29^ and kinesin-1^27^. As in Ref. 29, we use an event-based picture of motor dynamics, while including additional chemical and structural details specific to myosin-V-like motors (in particular the polymer properties of the lever arms). Similar to these earlier studies, we derive analytical expressions for observables like the distance, time, and velocity of runs, as well as their corresponding distributions^10, 27, 29^. The exact results often involve infinite sums and special functions that limit their practical usefulness, but we introduce an approximate treatment of the run velocity that greatly facilitates interpretation and analysis. This allows us to identify a key parameter, *α*, that assumes a value of 1 for motors with average run distances longer than one step or 2 for motors with average run distances less than one step.

For the case when *α* = 1, the mean velocity expression simplifies to the mean run distance divided by the mean run time (Eq. 25). Many theoretical studies of processive motors have used this definition of mean run velocity to great success^10, 20, 21, 29^, and we can now rigorously show in what context it is valid. This result may also apply to non-myosin-V-like processive motors so long as they translate along a periodic track and their motion partitions into forward and backward displacing events of uniform size and non-displacing events.

Continuing with our analysis of the mean run velocity, we described a Pareto frontier that defines the maximum average velocity obtainable by a motor with a given mean run distance (Eq. 29). This Pareto frontier offers insight into the design principles that govern the dynamic properties of a wide range of myosin-V-like motors. For processive motors with *α* = 1, the Pareto frontier assumes an inverse relationship between run distance and maximum run velocity. The tradeoff arises since increasing *k*_*d*1_ decreases dwell time while also increasing the termination probability. This region of the Pareto frontier offers theoretical validation of the experimental observation that fast motors sacrifice run distance for speed and vice versa^13^.

The right side of the Pareto frontier also offers a theoretical basis for the related observation that fast motors generate less force in a compliant optical trap (i.e. reach a smaller maximum trap displacement) than their slower counterparts^13^. Two model features impact maximum force generation: the stall force, which sets the maximum force obtainable by the motor, and the run distance, which determines to what extent the motor can reach its stall force. Based on these two factors, a motor’s force-generating behavior in a compliant optical trap may fall into two categories. The first is when the average run distance is sufficiently large to reach the stall force. In these cases, optical trap traces plateau before detachment, as demonstrated by *Nt*. myosin-XI in low ATP concentrations and myosin-V^14, 30^. Alternatively, the run distance may be shorter than the stall-force displacement, resulting in motor detachment before any observable stalling behavior. As noted by others, this second case likely explains the sharply peaked optical trap traces of *Nt*. myosin-XI at higher ATP concentrations^14^. Thus, as a motor along the Pareto frontier trades run distance for velocity, it also sacrifices its ability to generate force. One caveat to the above discussion is that the average run distance and velocity found in the Pareto frontier assumes a constant force while forces vary in compliant optical traps. However, the qualitative results likely hold since numerical simulations show that constant force run distance and velocity strongly correlate with compliant optical trap values (Fig. S4).

As the average run distance drops below one, representing *α* = 2 motors, the Pareto frontier assumes a constant value. This value is a global speed-limit set by the diffusive search of the free motor head for the forward actin binding site. Using the structural parameters for myosin-V, the speed-limit is approximately 35 *µ*m/s under no load. However, reaching this theoretical speed-limit requires perfect gating and infinitely fast enzymatic kinetics, which are bound by chemical diffusion rates (maximum of 10^8^ s^−1^) and enzymatic catalysis (maximum of 10^6^ s^−1^) in reality^31^. While the speed-limit may be an interesting theoretical feature, perhaps a more helpful finding is the square root relationship describing the *k*_*d*1_ rate that maximizes average run velocity at a given *k*_*h*_ value (Fig. 5). Using this and *k*_*h*_ = 10^6^ s^−1^ with myosin-V structural parameters, our model predicts that 22 *µ*m/s may be a more realistic unloaded speed-limit.

Despite our model’s speed-limit predictions, several studies of the ultra-fast *Cc*. myosin-XI report velocities of 60-70 *µ*m/s^15, 32^. However, these velocities were obtained from in vitro motility assays where ensembles of myosin motor domains fixed to a coverslip cooperatively translate actin filaments. A key theoretical study showed that mechanical coupling of multiple myosin by actin results in significant deviations from their single-motor dynamics where sliding velocity can increase several fold depending on the coupling strength^33^.

Our model’s predictions would therefore be ideally tested against optical trap studies of single motors traversing along actin filaments. Several experimental realities of such optical trap configurations will likely make this challenging. These include limited rise times and noise filtering during data post-processing, both of which attenuate the fast-changing signals produced by fast motors^14^. Quantifying non-productive runs poses a more fundamental issue since they are indistinguishable from Brownian noise yet significantly contribute to the average quantities predicted by our theory. One approach to overcome these experimental limitations is to fit a theoretical model to in vitro motility data. The fit could then be integrated with other kinetic and structural data to infer the single motor values of ⟨*Z*⟩, ⟨*τ*_*d*_⟩, and ⟨*V*⟩ using the expressions described here. While others have developed models of the in vitro motility assay that could be combined with our results, more theoretical and experimental work would be needed to accurately extract single-motor dynamics from ensemble behavior^33, 34^.

## Conclusion

To summarize, myosin-V-like molecular motors proteins have long been a subject of biophysical study^10^. Here, we propose a single theoretical framework that unifies structurally similar myosins with vastly different dynamics. In doing so, we offer expressions and limits that predict and constrain motor dynamics on the entire design space of myosin-V-like motors. We explicitly show that myosin-V-like motors make a tradeoff between average run distance and run velocity to function as drag racers or long haulers. However, myosin-V-like motors beyond myosin-V require significant structural and dynamic characterization to validate our predictions. The most important challenge to overcome will be to develop an approach that accurately assesses fast motor dynamics and non-productive behavior of single motors. Testing the predictions we make here will be foundational to our understanding of the unifying design principles governing molecular motor function.

## Materials and Methods

### Monte Carlo simulation

Code for this work was implemented using python v3.11 and accelerated with the numba, numpy, and packages. Exact package versions may be found in the “requierments.txt” file on the GitHub repository. Simulations were run on a slurm-based high-performance computing cluster. Analysis was performed using an interactive python environment in jupyter lab. Mathematica v12.0 was used as an aid in some analytical calculations. All code is maintained and publicly available on GitHub at https://github.com/hincz-lab/myo5space.

Two algorithmically distinct Monte Carlo techniques were used to simulate motor protein dynamics on the kinetic network found in Fig. 1. Both were used to assess the accuracy of each other’s implementation and to check the validity of analytical results. The first is a conventional rejection free kinetic Monte Carlo (kmc) simulation that randomly determines which connected state the motor transitions to, given its current position and appropriate transition rates. During the simulation, the current displacement of the motor and the total time it spends on the network are recorded.

The second technique takes advantage of the event-based picture of motor dynamics and is dubbed event Monte Carlo (emc). During the emc, an event from Tab. 1 is randomly chosen based on the calculated probability of each event happening. The consequential displacement change is recorded, and the event time is added to the overall run time. Similarly to the kmc, the emc continues until the terminal condition is met.

The nature of the terminal condition in both the kmc and emc depends on the quantity of interest. For simulations intended to calculate run distances, run times, and run velocities, the terminal condition is transitioning to a detached state or drawing a termination event in the case of the kmc or emc, respectively. If the goal is to record dwell times, the terminal condition is a displacing/terminating state transition or event. Finally, the simulations can simulate a motor working against a constant or Hookian resistive load according to the following equation.

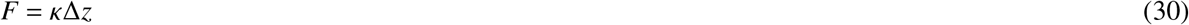

The latter represents the compliant optical trap used to study *Nt*. myosin-XI, while the former is helpful for other model evaluations. In the case of a compliant trap, the model’s force-dependent parameters, 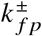, are updated following each simulation loop according to the motor’s current position.

### Model fitting

To fit the model to experimental data^14^, we use Monte Carlo simulations with a zero-order Bayesian maximum a posteriori approach. A zero-order approach was chosen since the fitting landscape was ill-suited to higher-order fitting algorithms. As discussed in the main text, we use distributions of dwell times and detachment forces collected at 1 *µ*M and 100 *µ*M ATP, substituting the dwell time distribution at 100 *µ*M ATP with a Gaussian prior on the mean run velocity,

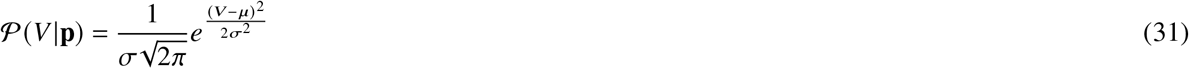

where *V* is the model’s predicted mean run velocity given a set of parameters **p**, *µ* (4.27 *µ*m/s) is the experimentally determined mean run velocity with standard error of the mean *σ* (0.35 *µ*m/s).

Fifty thousand trials of the appropriate simulations were run to generate distributions over a predefined grid of model parameters. All simulations were run in variable force mode to mimic the compliant optical trap used in experiments. The authors report a trap stiffness of 4.6 fN/nm and a bead-myosin linkage stiffness between 21 and 140 fN/nm. Assuming that these stiffnesses act in series, the overall stiffness of the experimental setup ranges between 3.77 fN/nm and 4.45 fN/nm. We thus use the mean stiffness of 4.1 fN/nm for simulations.

Before calculating the posteriors, several preprocessing steps were performed. First, several thresholds and cutoffs were enforced to mimic the inclusion criteria used by the experimental study^14^. Only runs that reached a minimum force threshold of 0.25 pN were used for detachment data, while a threshold of 0.5 pN was used for dwell time data. Due to the temporal resolution of the experimental apparatus used, the authors also imposed a 20 ms threshold for dwell time samples. For average run velocity calculations, the authors only used data below 0.3 pN so to only study speeds at low force. We further required that at least 1% of the original samples remain after enforcing the cutoffs to avoid instances where less than 500 data points are used for creating the simulated distribution. Distributions that failed to meet this final threshold were not considered viable parameter choices.

Next, the simulated data were converted into probability density functions (pdf). A Gaussian kernel density estimator using Scott’s rule for bandwidth estimation was used for dwell time distributions. The discreet distribution was first generated from the simulation data for detachment distributions. Then a pdf was generated using a sum of Gaussians with means equal to the detachment values and weighted by the probability of those detachment values. The standard deviation of each Gaussian was taken as 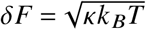 which describes the fluctuations in force data for a compliant optical trap with stiffness *κ* and at temperature *T* ^35^. Lastly, these distributions were integrated over each bin used to generate the experimental detachment and dwell time distributions. The resulting probabilities were used to calculate the natural logarithm of the likely hood or prior for each parameter set for a given experiment. The four individual fitting results were combined by first adding the results for a given parameter set at the same ATP concentration. The results at different ATP concentrations were further combined to allow *k*_*d*1_ to assume a different value at each ATP concentration. Finally, the parameter set with the maximum log posterior was selected as the best fit.

## Acknowledgments

This work made use of the High Performance Computing Resource in the Core Facility for Advanced Research Computing at Case Western Reserve University. We also would like to thank the authors of the experimental study of *Nt*. myosin-XI (Kazuhiro Oiwa, Motok Tominaga, and Hiroaki Kojima, ref 14), for a very insightful correspondence explaining the details of their experimental setup.

## Author contributions

**Joshua Holmes** Software, Formal analysis, Validation, Investigation, Visualization, Methodology.

**Michael Hinczewski** Conceptualization, Supervision, Investigation, Visualization, Methodology, Project administration.

## Supplementary materials for

### Drag racers and long haulers: the design space of myosin-V-like motor proteins

#### Calculating mean first passage rates

Here we catalog the equations used to calculate the mean first passage rates, 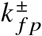 (Eq. 1 of the main text). Full derivations and justifications for any approximations used can be found in Ref. 1.

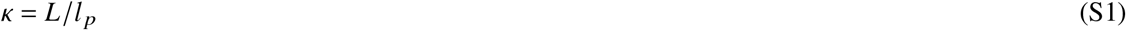

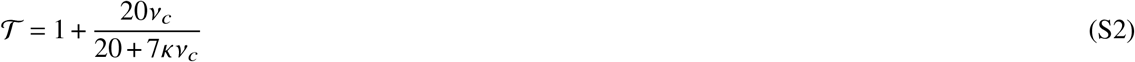

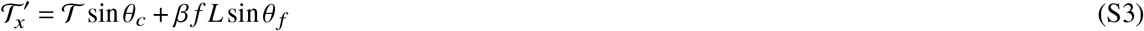

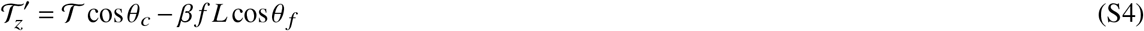

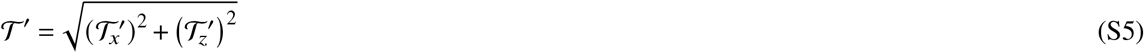

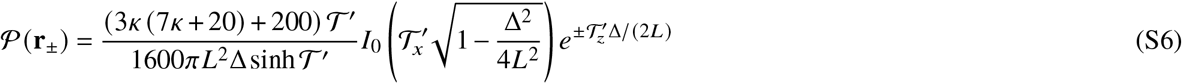

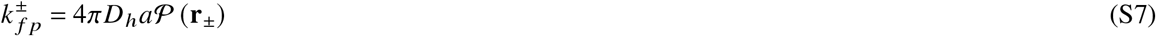

The parameters Δ, *L, l*_*p*_, *θ*_*c*_, *v*_*c*_, *f*, *θ*_*f*_ are as defined in the main text.

#### Event Probabilities

The expressions for ℙ_*i∈E*_ as defined in the main text are as follows.

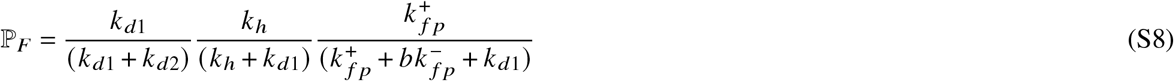

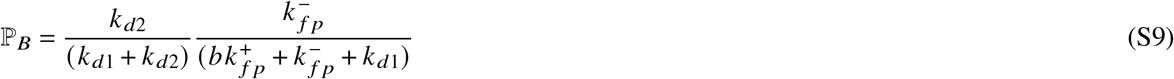

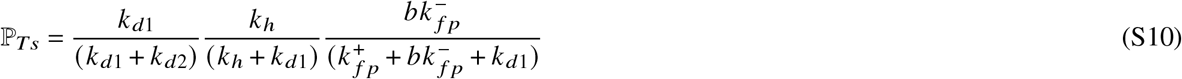

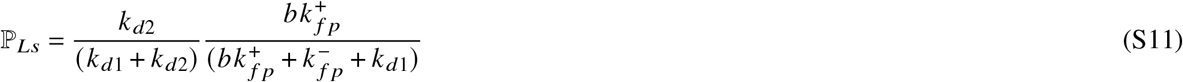

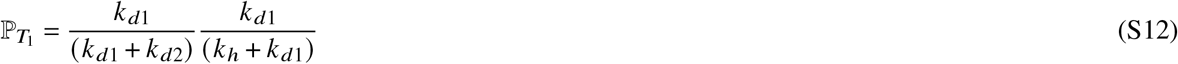

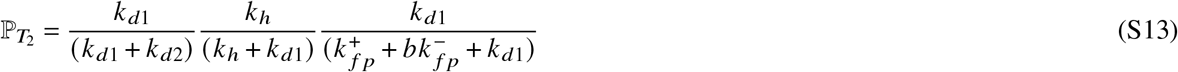

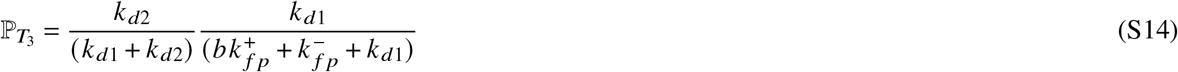

#### Justifying excluding the experimental dwell time distribution at 100 *µ*M ATP

We found that the reported distribution of dwell times at 100 *µ*M ATP was inconsistent with the measured average run velocity and led to overall poor fitting results. Dividing the motor’s mean step size of 35 nm by the experimental average dwell time at 100 *µ*M ATP of 25 ms would predict a mean run velocity of 1.4 *µ*m/s, which is significantly lower than the reported motor velocity of 4.27 *µ*m/s. This led us to instead incorporate the experimental run velocity information at 100 *µ*M ATP with a Gaussian prior parameterized by the reported mean and standard error.

### Derivation of the run distance distribution

Here we derive the expressions for the run distance distribution in terms of the Gaussian hypergeometric function,

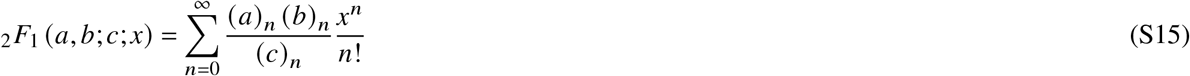

where (*a*)_*n*_ = *a* · (*a* + 1) · · · (*a* + *n* − 1) is the Pocchammer symbol with argument *a* and parameter *n*. The Pocchammer symbol can also be expressed in terms of the Gamma function 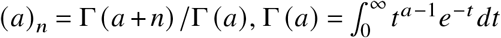.

First consider that the probability of a trajectory with *i* forward steps and *j* backwards steps is,

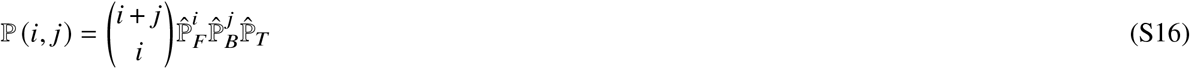

where the factor in the parentheses is the binomial coefficient, which accounts for every possible sequence of *i* and *j* forward and backwards stepping events. The final factor accounts for the obligatory termination event at the end of every run. Every trajectory where *z* = *i* − *j* contributes to the overall probability that the motor runs *z* steps,

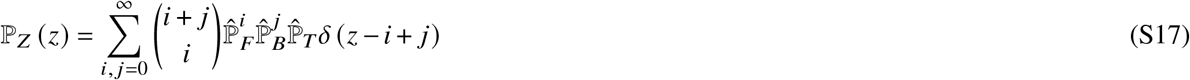

where *δ* is the Kronecker delta function. If we first consider the case where *z* ≥ 0, completing the sum over *i* produces,

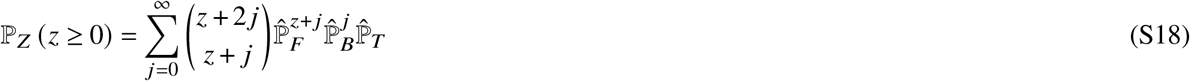

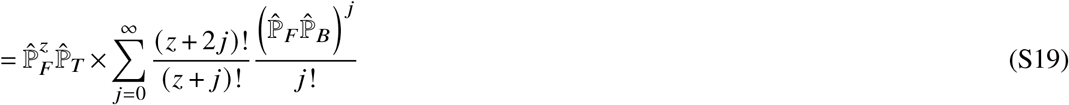

Now we manipulate the quotient of factorials.

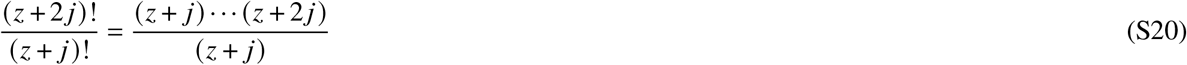

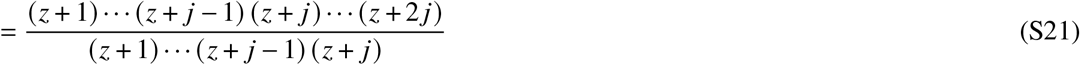

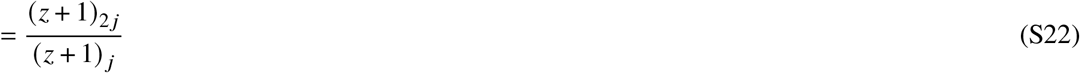

To proceed, we have to get the Pochhammer symbol in the numerator of in terms of Pochhammer symbols with parameter *j*. Using the duplication formula for the Gamma function as defined in Eq. 6.1.18 of Ref. 2,

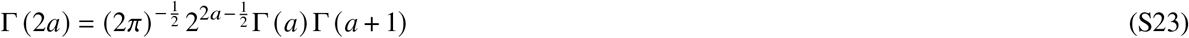

one can derive the following relationship.

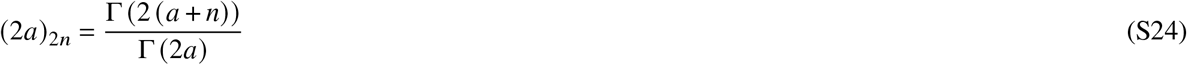

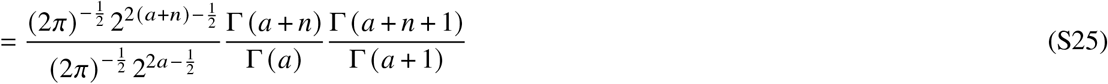

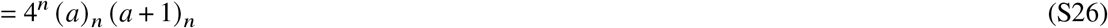

Now, the equation can be rewritten as,

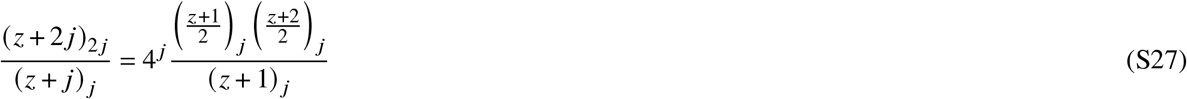

Substituting the above into Eq. S19 we arrive at the hypergeometric form of the run distance distribution.

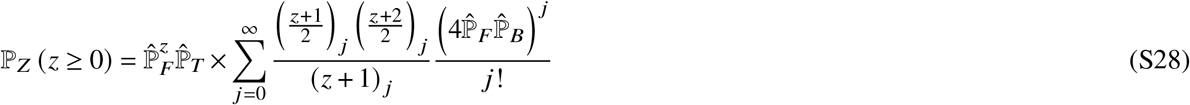

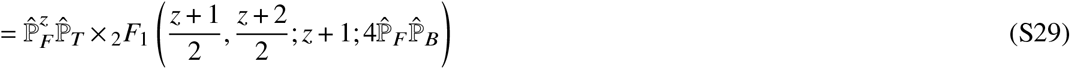

As mentioned in the main text, the hypergeometric function has the form,_2_(*F*_1_ *a*; 1 /2 + *a*; 2*a*; *x*) and has a closed form representation (Eq. 15.1.14 in Ref. 2),

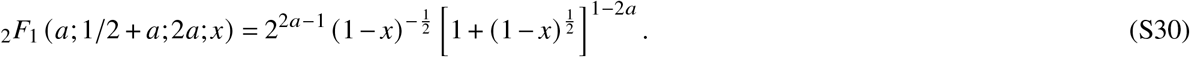

Using *a* = (*z* + 1) /2 gets the run distance distribution into the form reported in the main text,

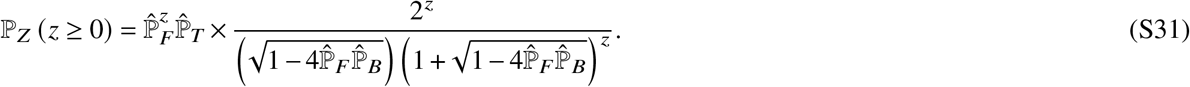

The case for *z* < 0 occurs when there are more backward steps than forward steps. Mathematically, this is equivalent to enforcing the condition that *z* = *i* − *j* < 0. Doing so switches the roles of *i* and *j* in Eq. S19 and the rest of the analysis continues as it was for the *z* ≥ 0 case. The end result is that *F* and *B* are interchanged with each other. For compactness, we introduce an indicator function, Φ(*z* ≥ 0) = *F*, Φ(*z* < 0) = *B*, to indicate directionality,

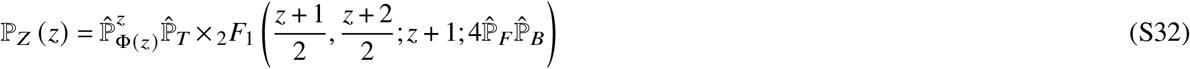

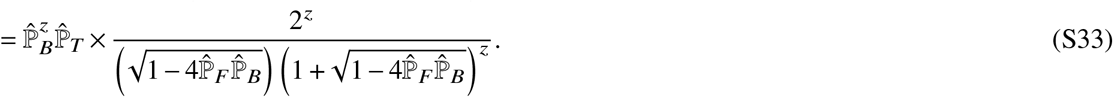

#### Derivation of the average dwell time

Here we derive the expression for the average dwell time. The average from main text Eq. 9 is,

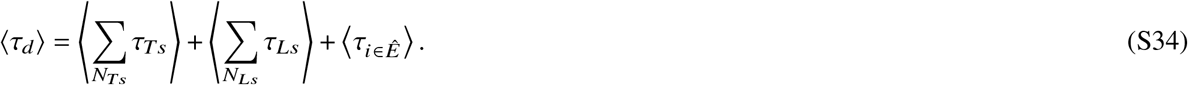

To manipulate the expression into something more transparent, note that the average operator distributes to each term in the two sums. In general, the average of a sum of *N* i.i.d. random variables, *τ*, is *N* ⟨*τ*⟩. If *N* is a random variable, then the average is ⟨*N* ⟨*τ*⟩⟩ = ⟨*N*⟩ ⟨*τ*⟩ since the average of a random variable is no longer a random variable. On average, there will be ⟨*N*_*Ts*_ and ⟨*N*_*Ls*_⟩ trailing and leading stomping terms, respectively. Furthermore, the final term representing the average transition time is the sum of the average event-times in *Ê* weighted by the associated event-probabilities on *Ê*, 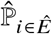 as defined in Eq. 3 of the main text. This can be shown formally by first noting that the probability density of *τ*_*i ∈ Ê*_ is a mixture of the densities of the members of *Ê*, 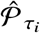, weighted by their respective probabilities 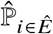.

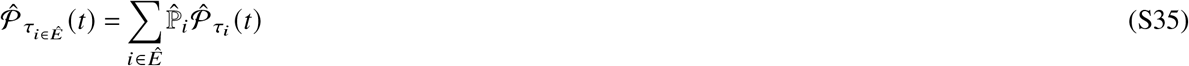

The rest follows from using the definition of an expectation value.

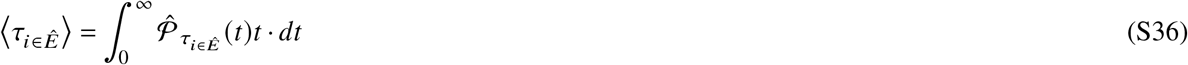

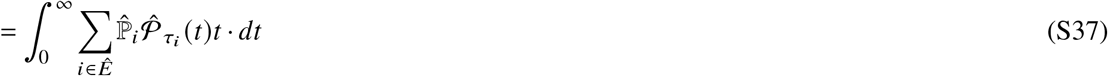

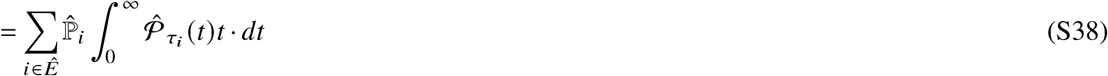

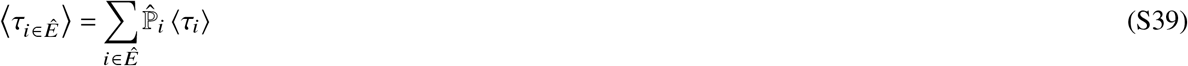

Exchanging the sum and integral is allowed since there are a finite number of terms in *Ê*. Now we have,

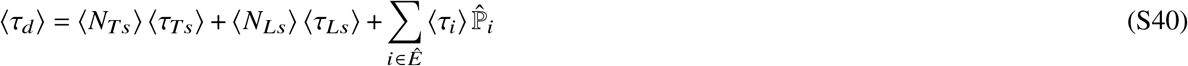

One calculates the average stomp numbers by partitioning the average total number of stomps 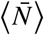 between trailing and leading stomps according the the relative probability of each. Since the leading and trailing stomp events constitute the set Ē, these relative probabilities are those of 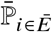 as defined in Eq. 4 of the main text. Thus,

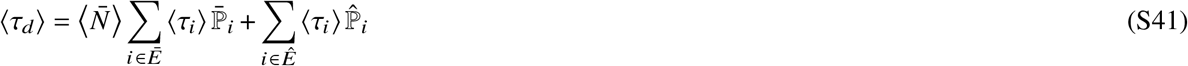

This is the expression reported in the main text.

As the main text also mentions, the average number of stomps before the transition event is geometrically distributed. For completeness, the specific form of the distribution is reported below.

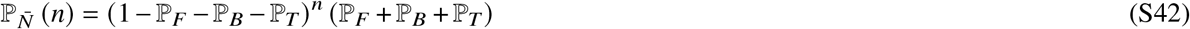

Its average is, 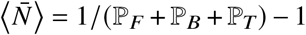, as identified in the main text.

The expressions for the average event-times ⟨*τ*_*i∈E*_⟩ in terms of the kinetic rates are as follows.

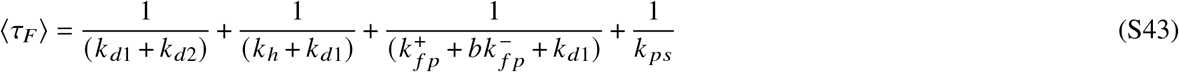

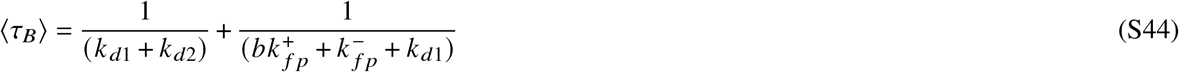

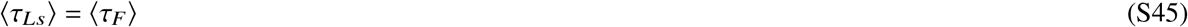

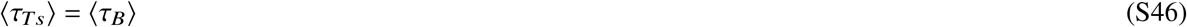

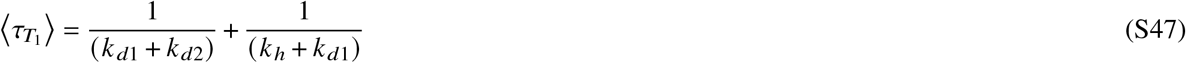

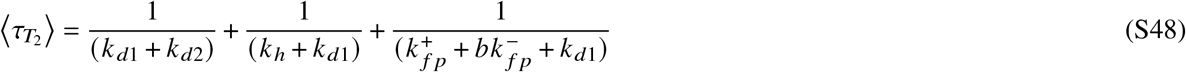

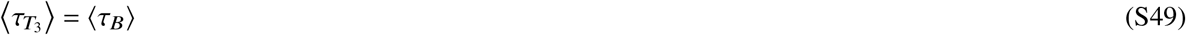

### Alternative expression for the average run time

As the main text mentions, we can express the run time in terms of a sum of event times, *τ*_*r*_ =_*N*_ ∑_*N*_*τ*_*i∈E*_. Following the same arguments used in deriving S41, the average run time in this cases becomes ⟨*τ*_*r*_ ⟩ = ⟨*N*⟩ ∑_*i∈E*_ ⟨*τ*_*i*_⟩ ℙ_*i*_. As shown in the main text, ⟨*N*⟩ = 1/ ℙ_*T*_. Thus,

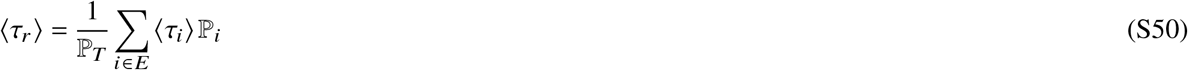

where the average event times and event probabilities are reported earlier in this SI.

### Dwell time and run time transition matrices

The transition matrix for representing a kinetic network that describes a motor’s dwell time at a single position on the actin track is,

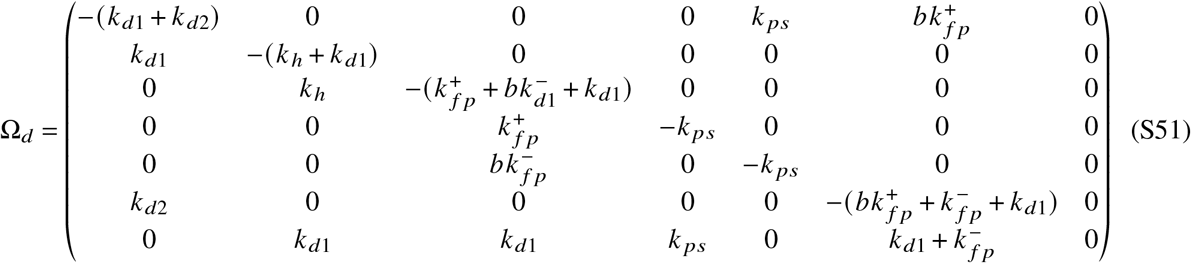

The index of each column corresponds to the index of the states in Fig. 1 of the main text (1: resting position, 2: pre-hydrolysis trailing head detached, 3: post-hydrolysis trailing head detached, 4: pre-powerstroke forward step, 5: pre-powerstroke trailing stomp, 6: leading head detached, 7: absorbing state). Column seven represents the absorbing state that replaces the final state in the sequences *S*_*i*∈*Ê*_, as defined in the main text.

The initial probability vector is a column vector such that the motor is guaranteed to start in the resting state (i.e. column one of Ω_*d*_).

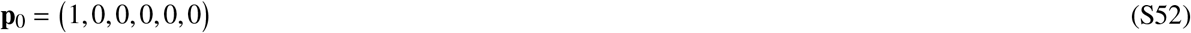

As mentioned in the main text, way to describe the distribution of run times is to use a network where only the terminating events connect to the absorbing state. In other words, the motor gathers time traversing the network while completing steeping and stomping events until both legs detach from actin. An advantage is of this picture is that the run time is no longer the sum of some other random variable. One simply needs the necessary transition matrix, which is essentially the same as S51 except for the bottom row to reflect the differences in absorbing conditions.

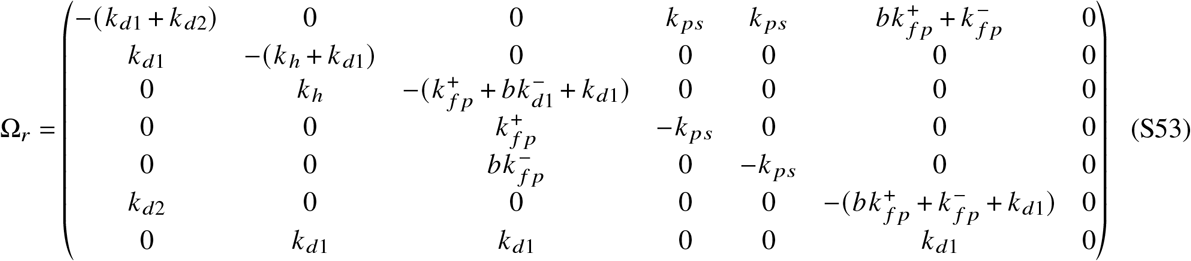

Ω_*r*_ has a submatrix *A*_*r*_ according to the equivariant of main text Eq. 14 and row vector **a**_*r*_ = −**1***A*_*r*_. The initial probability vector is the same as Eq. S52. Now, the run time distribution is,

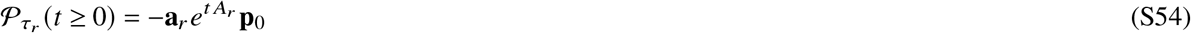

### Problems with the phase-type distribution

To illustrate the problem with the phase-type distribution, consider the distribution of a finite sum of 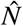 i.i.d. dwell times. Since the dwell times themselves are phase-type distributed, a closure property of phase-type distributions ensures that the sum will also follow a phase-type distribution with a new 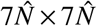 transition matrix, 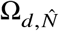^3^. To construct 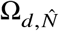, merely chain the absorbing state to the resting state for the appropriate amount of Ω_*d*_ matrices. The new probability vector begins as before, with the necessary number of zeros added to the end. Since 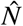 is geometrically distributed, the run time distribution will be an infinite mixture of geometrically weighted phase-type distributions.

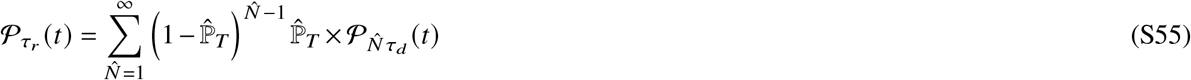

Here, however, lies a critical problem. While a second closure property further guarantees that the sum in Eq. S55 is also a phase-type distribution for a finite number of terms, it breaks down for infinite mixtures^3^. To see the source of the problem, consider how as 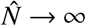, the dimension of 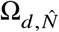 also becomes infinite. Clearly, matrices of infinite dimension are neither analytically practical nor amenable to computational approaches.

As a separate issue, the matrix equations used to describe the phase-type distribution offer little insight into the nature of the design space of myosin-V-like motors. For instance, even though main text Eq. 16 and main text Eq. 10 describe the same value, main text Eq. 10 offers a more transparent view of what motor properties contribute to the mean dwell time. Thus, we need an adequate approximation for the phase-type distribution of dwell times that offers both convenient analytical properties and transparency.

### Derivation of the average run velocity and its special case

Here we derive the general expression for the average run velocity (Eq. 24 of the main text) and its special form when the approximating dwell time shape parameter, *α* = 1. We start with the definition of an average.

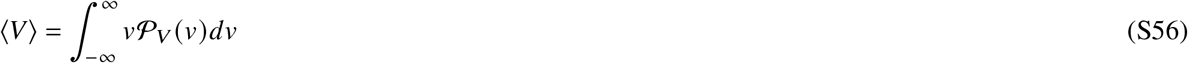

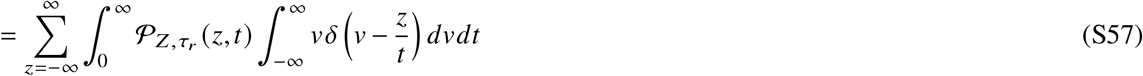

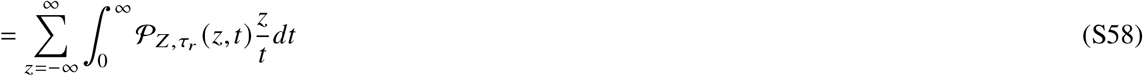

where *δ* is the Dirac delta function. Replacing the joint density with main text Eq. 23,

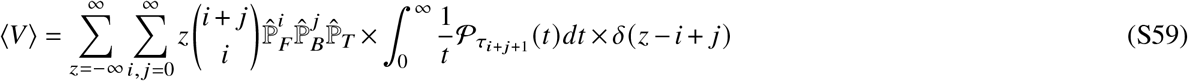

Before using the delta function to get rid of either *i* or *j*, it is useful to split the sum over *z* into its positive and negative halves. We can convert the sum over negative values of *z* into a sum over positive *z* values by replacing *z* with −*z*. In doing so, we also swap *i* and *j* since *Z* = *i* − *j* → −*Z* = *j* − *i*.

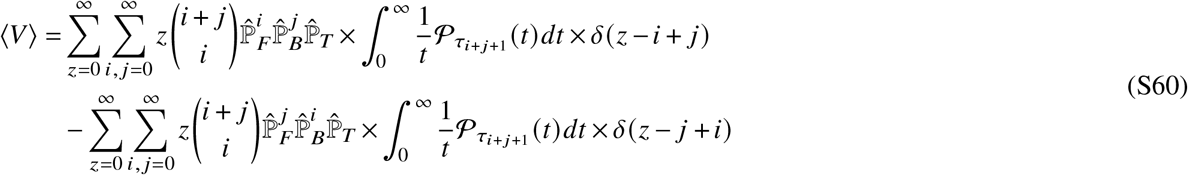

Note that the binomial coefficient and the density for *τ*_*i*+ *j*+1_ are symmetric in *i* and *j*. Now we can perform the sum over *i* in both terms.

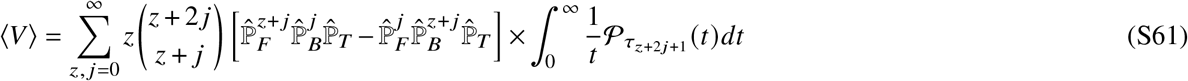

The integral quantity represents the first negative moment of the distribution of *τ*_*z*+2 *j*+1_. The first negative moment for a given distribution can be calculated using the distribution’s moment generating function, 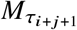 ^4^.

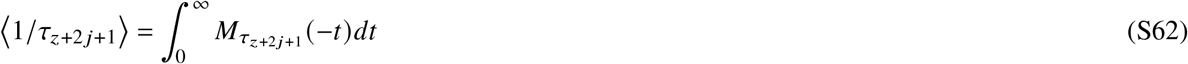

Since we are approximating *τ*_*z*+2 *j*+1_ as a gamma distribution, the relevant moment generating function is *M*_*τ*_*z*+2 *j*+1 = (1 − *θt*) ^−(*z*+2 *j*+1) *α*^. Making the necessary substitution into the above and performing the integration via a change of variables, we get an expression for the first negative moment.

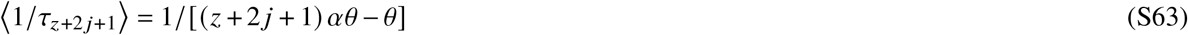

For a general gamma distribution, (*z* + 2*j* + 1)*α* is replaced with whatever the appropriate shape parameter is. Now the average run velocity is in terms of probabilities, approximating gamma parameters, and indices.

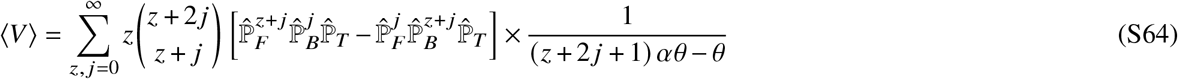

To get the average run velocity in terms of the hypergeometric function as reported in the main text, we first reformat the above to isolate the sum over *j* and express the binomial coefficient in terms of factorials.

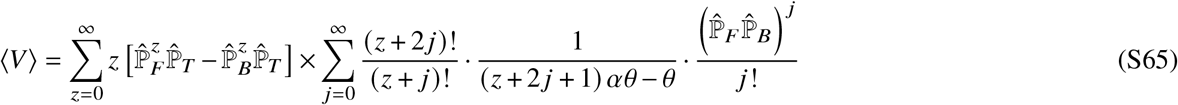

We can recast the quotient of factorials in terms of Pochhammer symbols parameterized by *j* using Eq. S22. The second factor in the sum over *j* can also be expressed in terms of Pochhammer symbols.

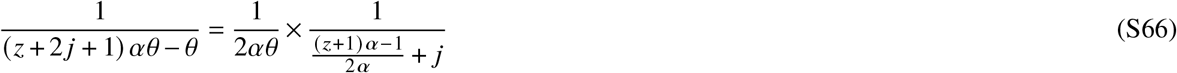

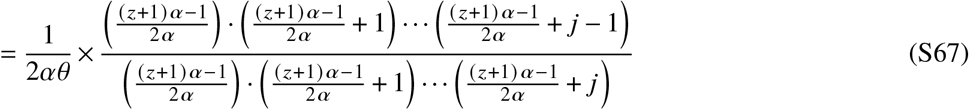

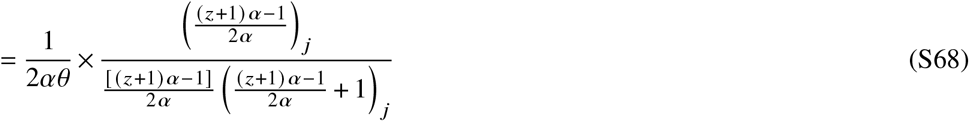

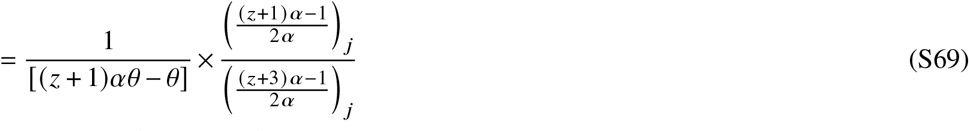

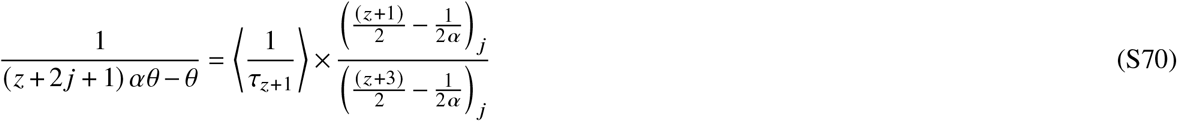

Using the results from Eq. S22 and Eq. S70 in Eq. S65 the sum over *j* now has the form of a generalized hypergeometric function.

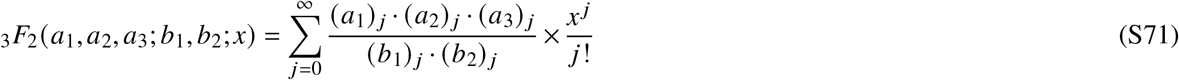

Now the average run velocity assumes the form reported in the main text.

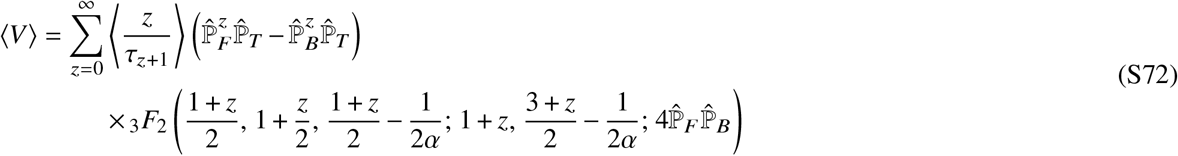

We also mention in the main text that the average run velocity has a special form when *α* = 1. Evaluating the above with *α* = 1 causes the second and fifth arguments to be the same. Since one is in the numerator and the other in the denominator, they cancel and reduce the Gaussian hypergeometric function to,_2_*F*_1_([*z* + 1]/2, [*z* + 2]/2; *z* + 1; 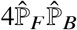).

This has the same form as in Eq. S30 with *a* = (*z* + 1) / 2. Furthermore, using Eq. S62, ⟨*z*/[*τ*_*z*+1_⟩|_*α*=1_ = 1/*θ*. Since 1/ *θ* = 1/ [*αθ*], we find that ⟨*z*/[*τ*_*z* + 1_]⟩|_*α*=1_ = 1/⟨*τ*_*d*_⟩. Plugging this into the above reduces it to a difference of geometric series.

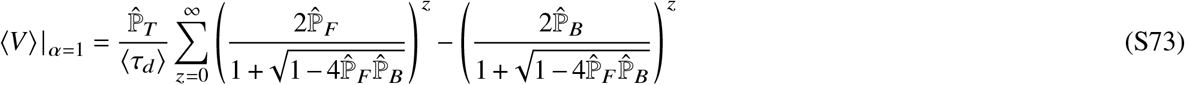

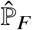 and 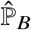 are bounded by zero and one, so both geometric series converge.

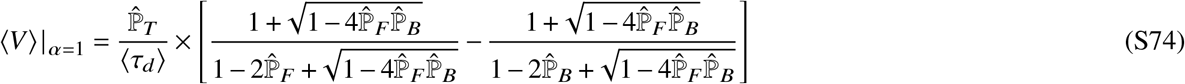

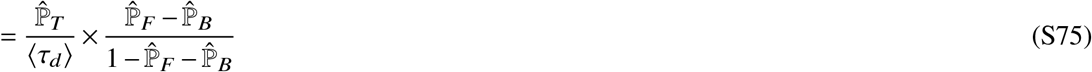

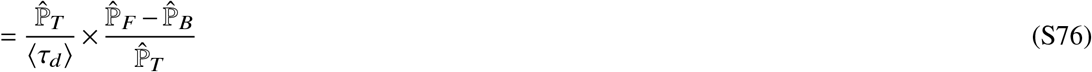

In this form, the first factor is the inverse of the average run time (main text Eq. 13) and the second is the average run distance (main text Eq. 5).

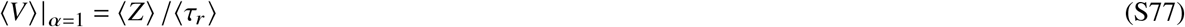

### Derivation of the myosin speed-limit expressions

Here we derive the various expressions used to find the myosin speed-limit.

First, the distribution of run velocities in the case of perfect gating looks is similar to the general run velocity distribution. The difference is in the distribution of run distances. Since a motor with perfect gating cannot take backwards steps, there is only one trajectory that can produce a given run distance: 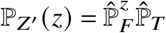. Note that we have used *Z*^′^ to denote the run distance when *k*_*d*2_ = 0. Much like before, the distribution of run times for these trajectories is the distribution of a sum of *z* + 1 dwell times. Using the gamma approximation for the dwell time distribution, 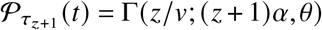, the joint density is then just the product of the run distance and run time distributions: 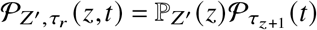. To get the run velocity distribution, just substitute this joint density with the one in Eq. 23 of the main text.

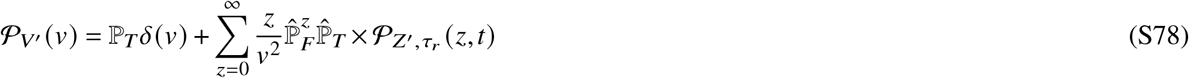

Note that the sum over *z* starts at zero since negative values of *z* are impossible.

To get the average run velocity with perfect gating, we start again with the definition of the average.

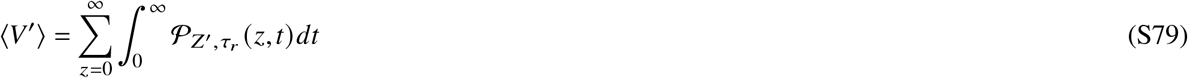

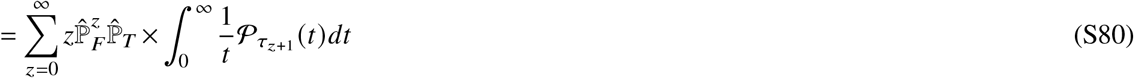

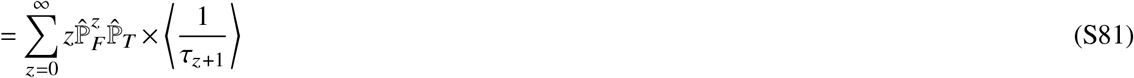

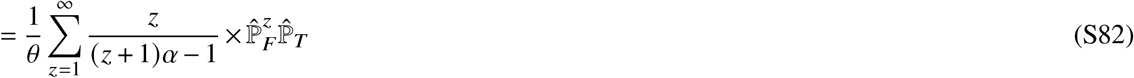

The last step was done by writing out the full expression for the negative-moment in terms of *α* and *θ* (Eq. S70). Also note the change in starting index in the last step since the fist term is zero. Now we can manipulate the first factor in the sum to get it in terms of Pocchammer symbols.

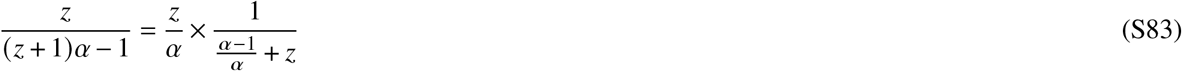

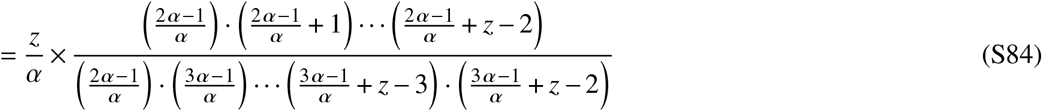

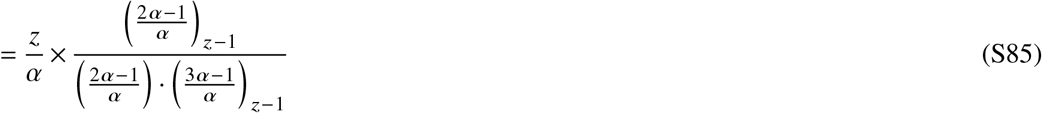

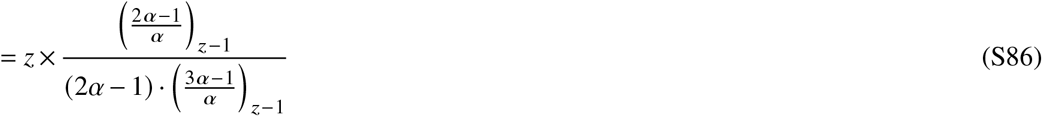

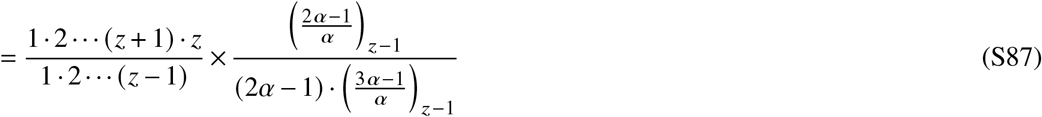

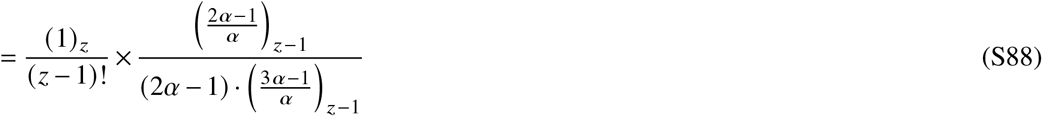

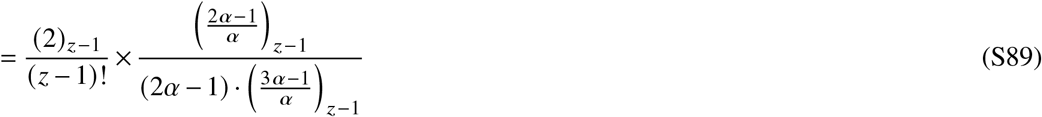

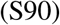

Replacing this into Eq. S82 allows us to simplify to a Gaussian hypergeometric function.

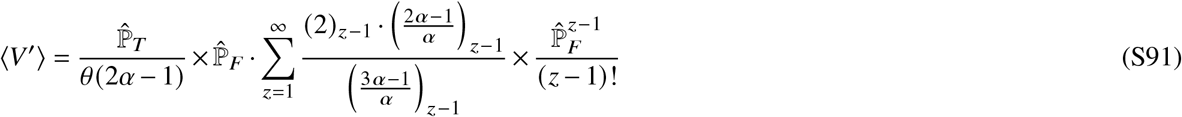

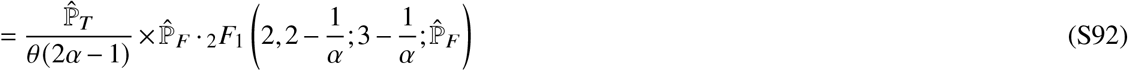

This form of the hypergeometric function is contiguous with another hypergeometric function,_2_*F*_1_ (*a, b*; *c, x*)∼_2_*F*_1_ (*a* + 1, *b* + 1; *c, x*), where *a* = 1, *b* = 1 − 1/*α, c* = 3 − 1 *α*, 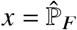. To find the exact mapping between the two, first use Eq. 63 from Ref. 5.

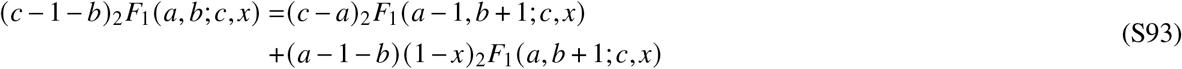

Since *a* = 1, the first argument of the first term is zero and that hypergeometric function is equal to one. Then, solving for_2_*F*_1_(*a, b*; *c, x*),

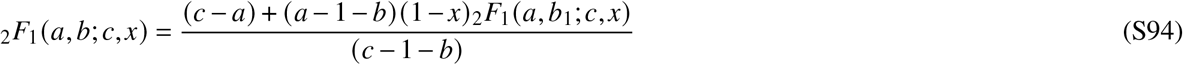

To get_2_*F*_1_(*a, b* + 1; *c, x*) in terms of_2_*F*_1_(*a* + 1, *b* + 1; *c, x*), use Eq. 42 from Ref. 5.

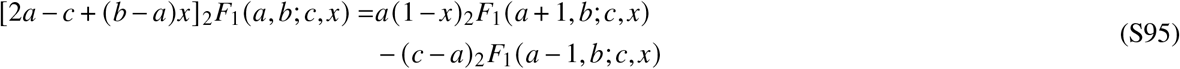

The hypergeometric function in the second term on the right-hand side again goes to one. Then, make the substitution *b* → *b* + 1 and solve for_2_*F*_1_(*a, b* + 1; *c, x*).

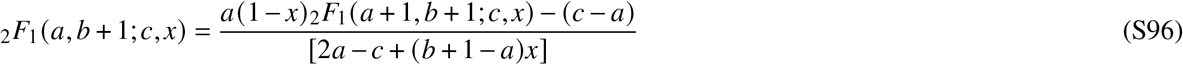

Now make the substitutions *a* → 1, *b* → 1 − 1/*α*, and *c* → *c* − 1/*α*.

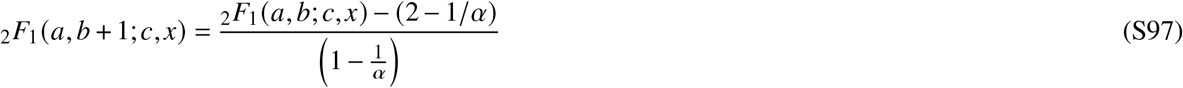

Replace the hypergeometric function in Eq. S94 with the above and make the parameter substitutions.

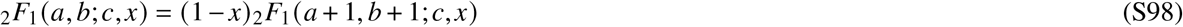

Finally, this can be used to simplify the expression in Eq. S92.

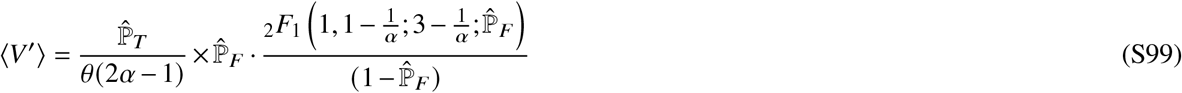

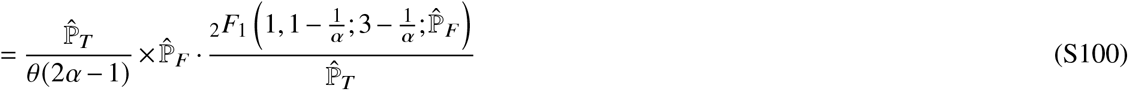

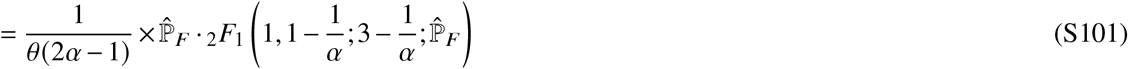

which is the form reported in the main text.

As mentioned in the main text, the above has a special form when *α* = 2. First evaluate the above with *α* = 2.

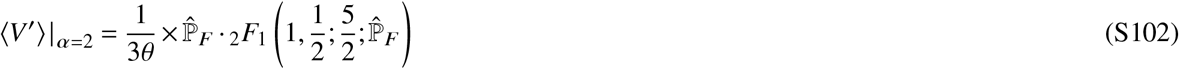

The hypergeometric function here is contiguous with_2_*F*_1_ (1, 1/2; 3/2; 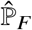) and share the following relationship as define by Eq. 15.2.27 of Ref. 2.

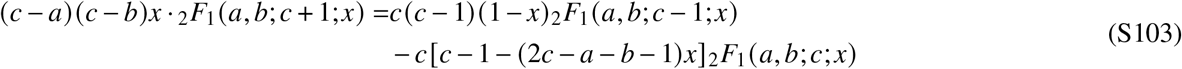

Now make the substitutions *a* → 1, *b* → 1/2, *c* → 3/2, and 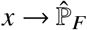.

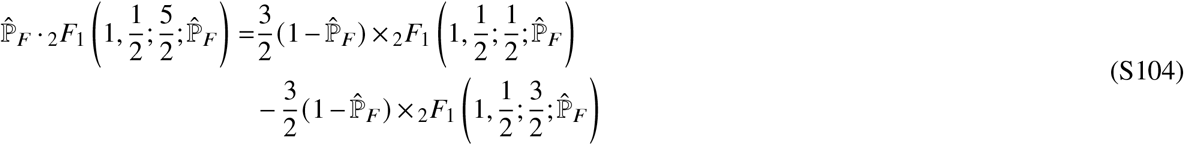

The hypergeometric function in the first term on the right hand side simplifies to 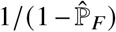 as in Eq. 15.1.8 of

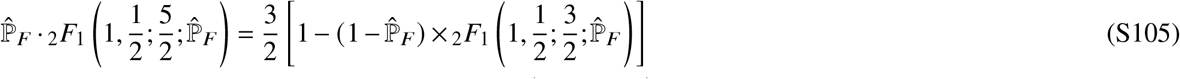

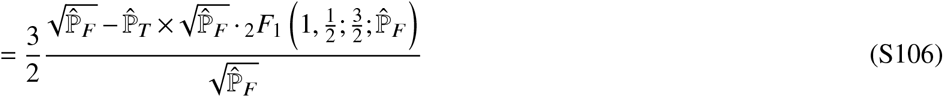

The final factor in the numerator is of the form *x*_2_*F*_1_ (1, 1/2; 3/2; *x*^2^) and has a closed form as defined by Eq. 15.1.4 of Ref. 2.

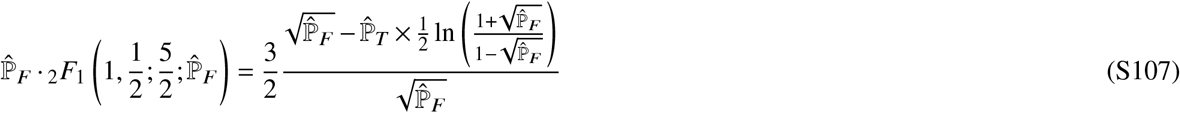

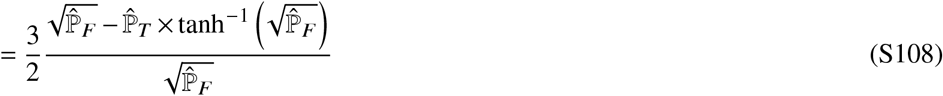

Replacing this into Eq. S102 gets the form reported in the main text.

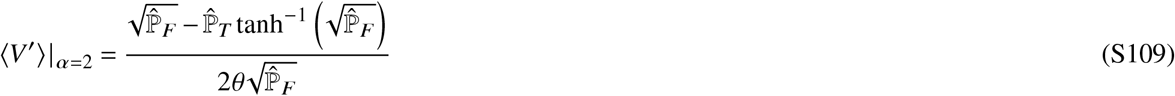

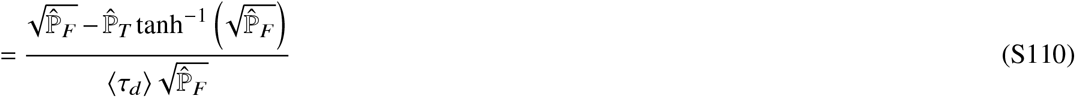

The final step is possible since 2*θ* = *αθ* = ⟨*τ*_*d*_⟩.

### Deriving the myosin Pareto-frontier

Here we derive the Pareto frontier that bounds the scatter plot of average run velocity versus average run distance (main text Fig. 7). Ultimately, the goal is to find an expression for the maximum average run velocity obtainable at a given average run distance.

We start with the region where ⟨*Z*⟩ > 1 and *α* ≈ 1. Much like in the derivation of the global maximum average run velocity, we can assume that motors at the Pareto-frontier will have perfect gating (i.e. *k*_*d*2_ = 0) and operate under constant force. Thus, we can evaluate ⟨*V* ^′^⟩ (Eq. S102) for *α* = 1 to get the run velocity at the frontier. In doing so, the second parameter in the hypergeometric function of Eq. S102 evaluates to zero so the hypergeometric function is simply one and the whole expression collapses to a simple quotient.

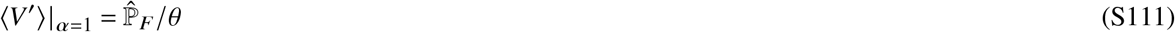

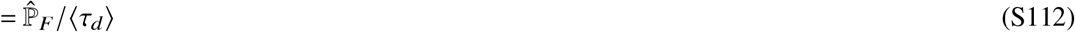

The last step is possible since *θ* = 1 · *θ* = *αθ* = ⟨*τ*_*d*_⟩.

In the limit of perfect gating, we can get 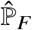 in terms of ⟨*Z*^′^⟩.

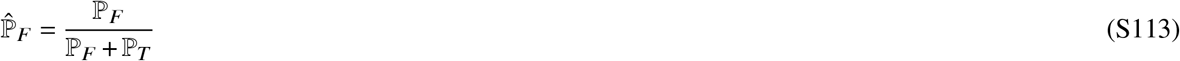

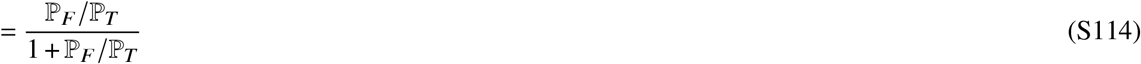

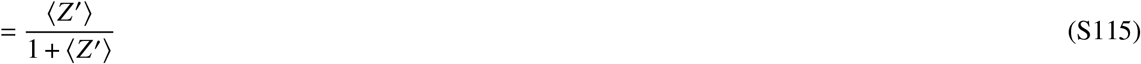

When ⟨*Z*⟩ ≫1, 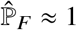 and ⟨*V* ^′^⟩|_*α*=1_ ≈ 1 / ⟨*τ*_*d*_⟩. Furthermore, consider main text Eq. 10 for the average dwell time. We just showed that 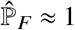, which means, by the normalization condition, that all the other probabilities are approximately zero. Therefore, in these limits, the average dwell time equals the average time it takes to complete a forward step: ⟨*τ*_*d*_⟩ ≈ ⟨ *τ*_*F*_⟩.

To proceed, we must work with Eq. S43 that relates ⟨*τ*_*F*_⟩ to the network’s rates. Following the argument that we make in the main text, we can make some simplifications by noting that *k*_*d*1_ will be the slowest rate for *α* ≈ 1 motors apart from *k*_*d*2_, which is zero in this case.

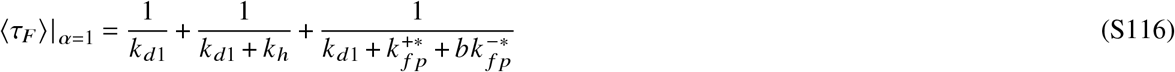

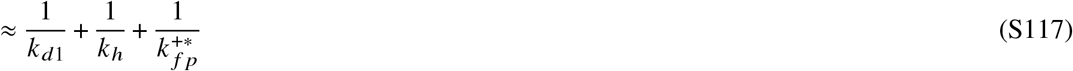

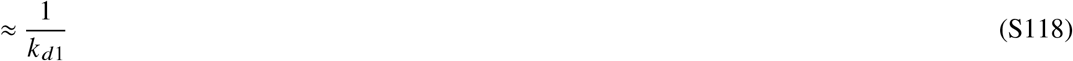

Note that we used 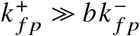 in the first step since ℙ_*F*_ ≫ ℙ_*Ts*_.

Using the work up until now, we can write the velocity at the frontier, ⟨*V*⟩^†^, for ⟨*Z*⟩ ≫ 1 as,

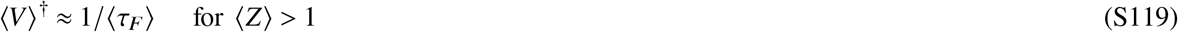

Next we need to get ⟨*τ*_*F*_⟩ in terms of ⟨*Z*^′^⟩. First we start with the expression for ⟨*Z*⟩ and approximate ℙ_*F*_ as one.

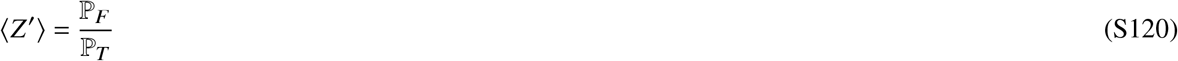

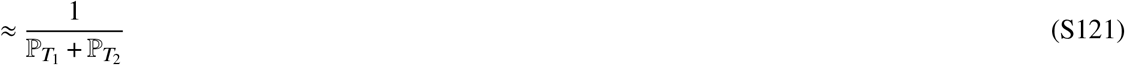

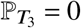 since *k*_*d*2_ = 0. We can use the fact that *k*_*d*1_ is the smallest non-zero rate for *α* ≈ 1 motors to simplify the denominator.

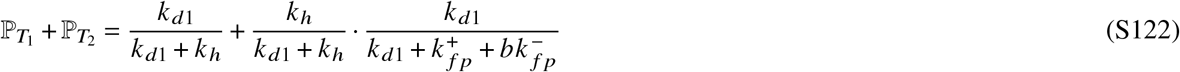

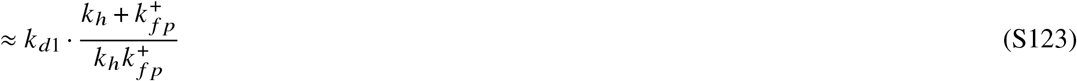

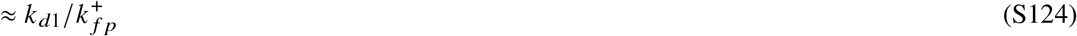

The final step was made by considering how the fastest motors at a given ⟨ *Z*^′^⟩ will have 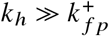. Replacing this into the above,

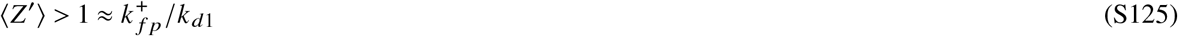

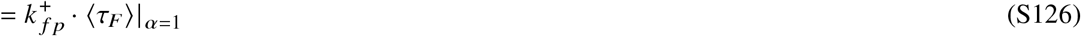

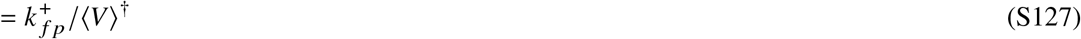

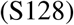

We obtain the bounding curve for the *α* = 1, or equivalently ⟨*Z*⟩ > 1, region by solving for ⟨*V*⟩^†^ and replacing ⟨*Z*^′^⟩ → ⟨*Z*⟩.

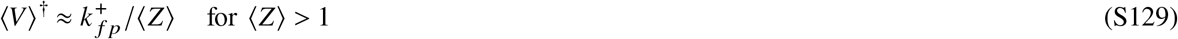

Now we move onto the case when ⟨*Z*^′^⟩ ≪ 1 and *α* ≈ 2. Since *α* ≈ 2, we can use ⟨*V* ^′^⟩|_*α*=2_ (Eq. S102) to calculate the average run velocities. As before, we need find the Pareto-frontier by getting Eq. S102 in terms of the average run distance under the assumptions that apply to the fastest motors. Unlike the ⟨*Z*^′^⟩ > 1 region, however, main text Fig. 7 shows that the frontier has two distinct parts. The first is relatively constant while the second is log-linear. If we consider that the log-linear boundary exists in the limit where ⟨*Z*^′^⟩ ≪ 1, then we can approximate 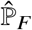 in Eq. S102 as ⟨*Z*^′^⟩.

The last job then is to simplify ⟨*τ*_*d*_⟩. To do so, we use the arguments made in the main text to note that *k*_*d*1_ with be significantly larger than any other rate in the network for motors with *α* ≈ 2 and ⟨*Z*^′^⟩ ≪ 1. I n this limit, 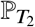 (Eq. S13) goes to one. Thus, the average dwell time (Eq. 10) will be approximately equal to 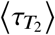. Furthermore, 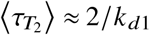. At a given ⟨*Z*^′^⟩ the fastest motor will have the largest 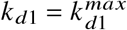. Since *k*_*d*1_ is a free parameter in our model, 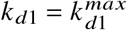 is technically infinity, but is finite for numerical evaluation. This would suggest that the log-linear boundary is an artifact. For completeness, however, we can still write down the equation for this fake frontier.

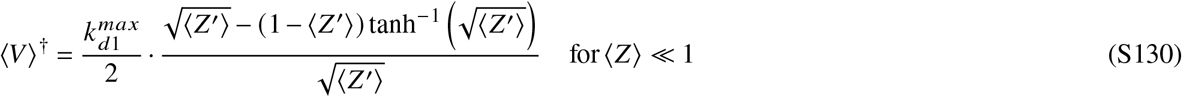

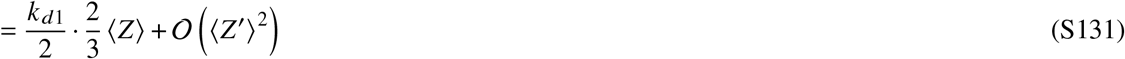

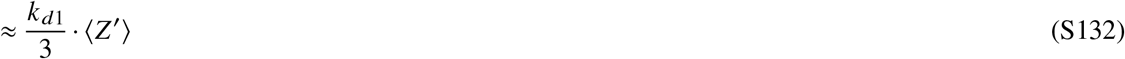

To rigorously prove that the log-linear frontier in the ⟨*Z*^′^⟩ ≪ 1 region is an artifact of a finite 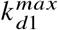, we can find an approximate ⟨*Z*^′^⟩ where the log-linear region begins. Based on the discussion in the main text, we use the ansatz that the transition between the constant and log-linear regions happens at 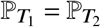. Since we are concerned with motors at the frontier, we can set 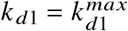, *k*_*d*2_ = 0, and 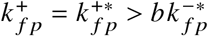, leaving *k*_*h*_ as the only unknown parameter. Thus, we can find the 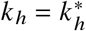 in terms of 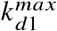 and 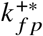.

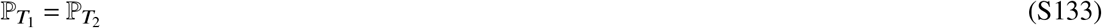

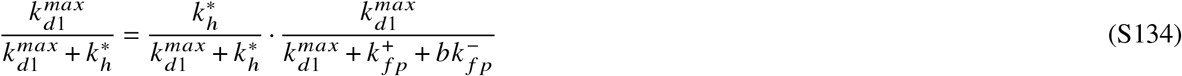

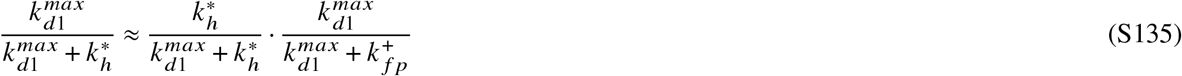

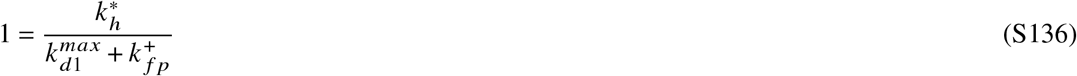

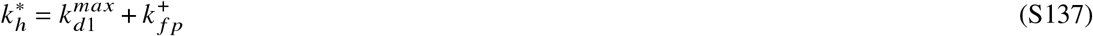

By evaluating Eq. S121 for ⟨*Z*^′^⟩ with 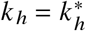 and using the usual approximations and assumptions, we can find an expression for the transition point.

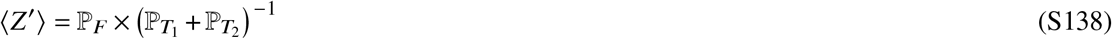

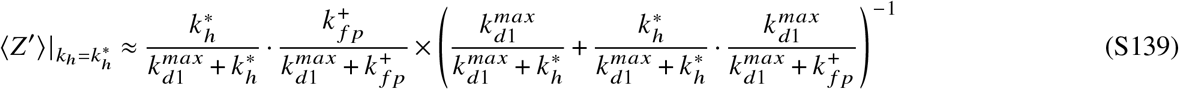

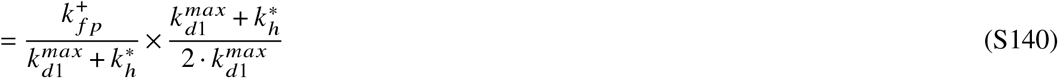

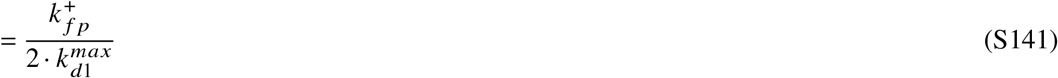

So, with 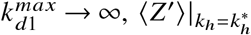 becomes arbitrarily close to zero. Therefore, the constant frontier actually accounts for the entire ⟨*Z*⟩ < 1 region. As one might expect, the value for the constant frontier is simply the maximum average run velocity defined in main text Eq. 28. Thus, the Pareto-frontier is,

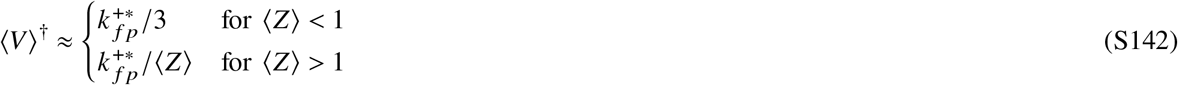

as reported in the main text.

Next, we show that the left asymptote of Fig. 7b is lim_⟨*Z*_′_⟩→ 0_ ⟨*Z*^′^|*Z*⟩ > 0 = 1. This asymptote represents the average run distance of productive runs of the fastest motors. We start by noting that the normalization condition on *Ê* is 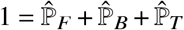. As in the derivation of the ultimate myosin-V-like speed-limit, we assumption that the fastest motors have perfect gating, implying 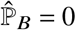. Thus, the normalization condition can be rewritten as 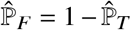. Another consequence of this perfect gating is that 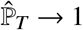 as ⟨*Z*^′^⟩ → 0 for motors at the speed-limit as can be seen from the definition of ⟨*Z*^′^⟩ (Eq. 5). We can then prove the asymptote is 1 by combining these with the definitions of ⟨*Z*^′^⟩ and ⟨*Z*^′^|*Z*^′^ > 0⟩.

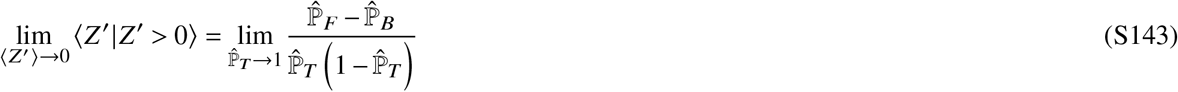

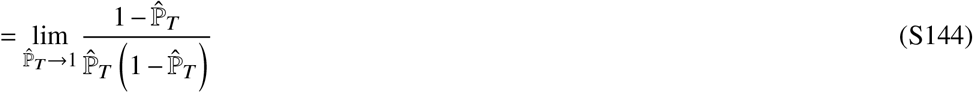

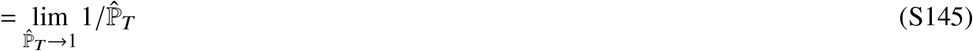

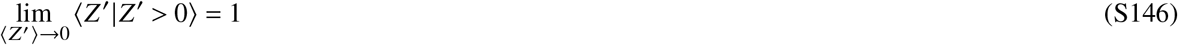

As we noted in the main text, motors with 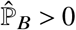(i.e. *k*_*d*2_ > 0) can break into the region to the left of the asymptote due to the effects of back stepping (Fig. S3b).

Finally, we show that the Pareto frontier bounding the right side of main text Fig. 7 does not change when the average run distance and run velocity are conditioned to productive runs: 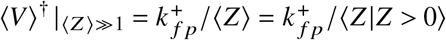. This hinges on showing that ⟨*Z*⟩ = ⟨*Z* |*Z* > 0⟩. We use the same assumptions as when originally deriving the rightward bounding Pareto frontier: 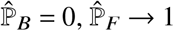, and ⟨*Z*^′^⟩ ≫ 1. As before, the normalization condition under these is 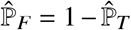. The derivation naturally follows.

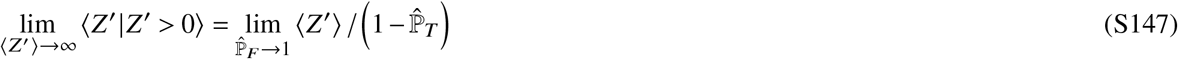

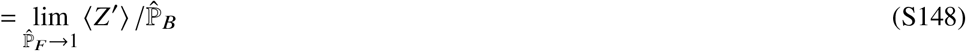

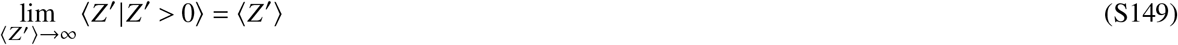

## Figures and Tables

**Table 1.** Summary of key analytical results. Reports the average and distributions, if present, for the run distance (*Z*), dwell time (*τ*_*d*_), run time (*τ*_*r*_), run velocity (*V*), the run velocity for motors with approximating shape parameters equal to one (*V*|_*α*=1_), and the Pareto frontier. Equation and symbol definitions may be found in the main text or SI where the equations are derived or discussed.

| Observable | Average | Distribution |
| --- | --- | --- |
| $Z$ | $(\hat{\mathbb{P}}_F - \hat{\mathbb{P}}_B) / \hat{\mathbb{P}}_T$ [Eq. 5] | $\frac{\hat{\mathbb{P}}_{\Phi(z)}^z \hat{\mathbb{P}}_T \times 2^z}{\left(\sqrt{1-4\hat{\mathbb{P}}_F\hat{\mathbb{P}}_B}\right)\left(1+\sqrt{1-4\hat{\mathbb{P}}_F\hat{\mathbb{P}}_B}\right)^z}$ [Eq. 8] |
| $\tau_d$ | $\langle \bar{N} \rangle \sum_{i \in \bar{E}} \langle \tau_i \rangle \bar{\mathbb{P}}_i + \sum_{i \in \hat{E}} \langle \tau_i \rangle \hat{\mathbb{P}}_i$ [Eq. 10] | $\mathbf{a}_d e^{t A_d} \mathbf{p}_0$ [Eq. 15] |
| $\tau_r$ | $\langle \tau_d \rangle / \hat{\mathbb{P}}_T$ [Eq. 13] | $\mathbf{a}_r e^{t A_r} \mathbf{p}_0$ [Eq. S55] |
| $V$ | $\sum_{z=0}^{\infty} \left\langle \frac{z}{\tau_{z+1}} \right\rangle \left( \hat{\mathbb{P}}_F^z \hat{\mathbb{P}}_T - \hat{\mathbb{P}}_B^z \hat{\mathbb{P}}_T \right) \times \Psi(z, \hat{\mathbb{P}}_F, \hat{\mathbb{P}}_B)$ [Eq. 24] | $\mathbb{P}_T \delta(v) + \sum_{z=-\infty}^{\infty} \frac{z}{v^2} \mathcal{P}_{Z, \tau_r} \left( z, \frac{z}{v} \right)$ [Eq. 22] |
| $V\vert _{\alpha=1}$ | $\langle Z \rangle / \langle \tau_r \rangle$ [Eq. 25] | |
| Pareto frontier | $\begin{cases} k_{fp}^+ / 3 & \text{for } \langle Z \rangle < 1 \\ k_{fp}^+ / \langle Z \rangle & \text{for } \langle Z \rangle > 1 \end{cases}$ [Eq. 29] | |

**Figure S1.**
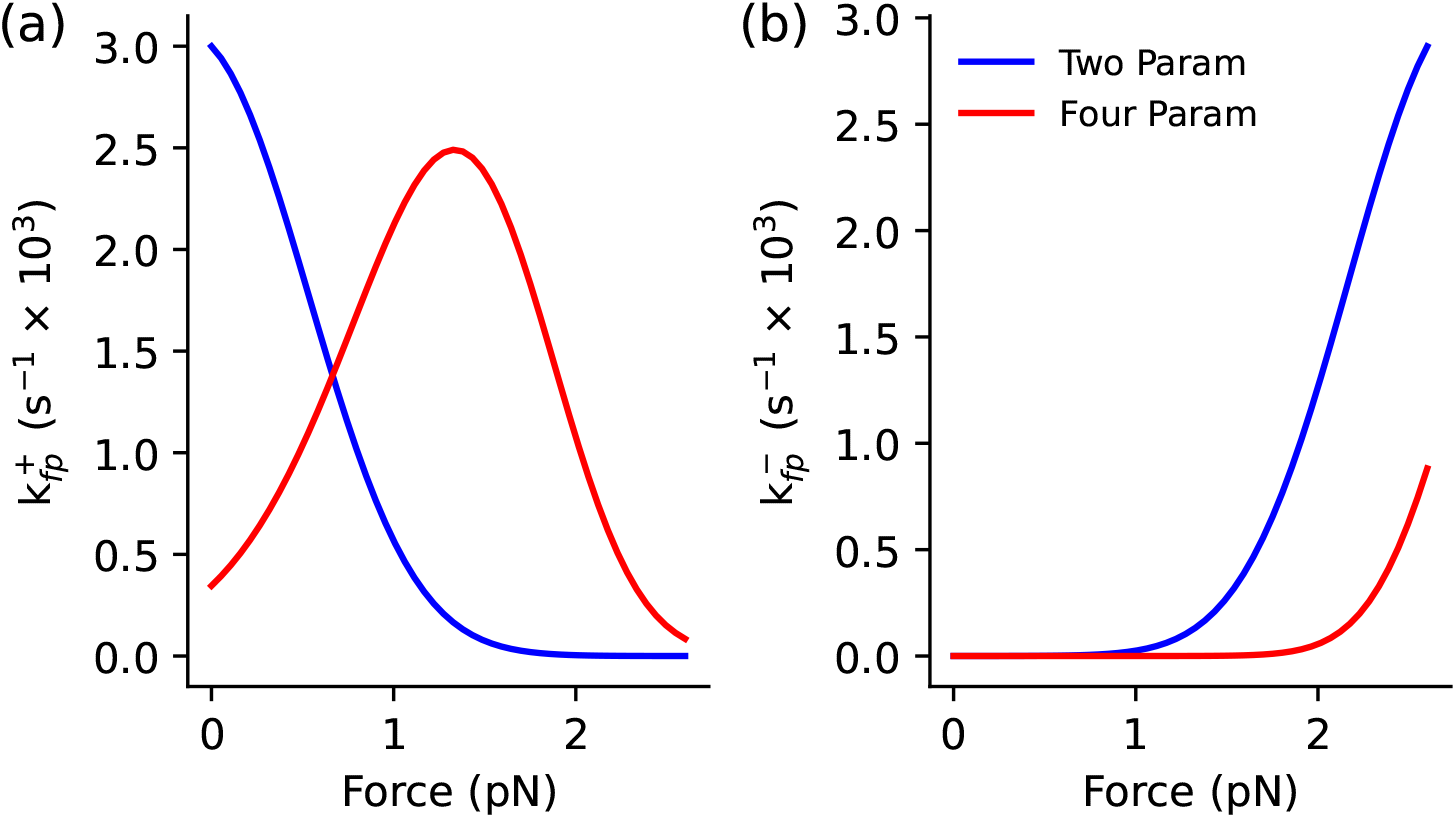
Force dependence of 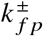 for motors fitted to *Nt*. myosin-XI data. The force dependence of (a) 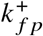 and (b) 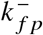 for motors parameterized according to the the two parameter fit (blue) and four parameter fit (red) as defined in main text Tab. 2.

**Figure S2.**
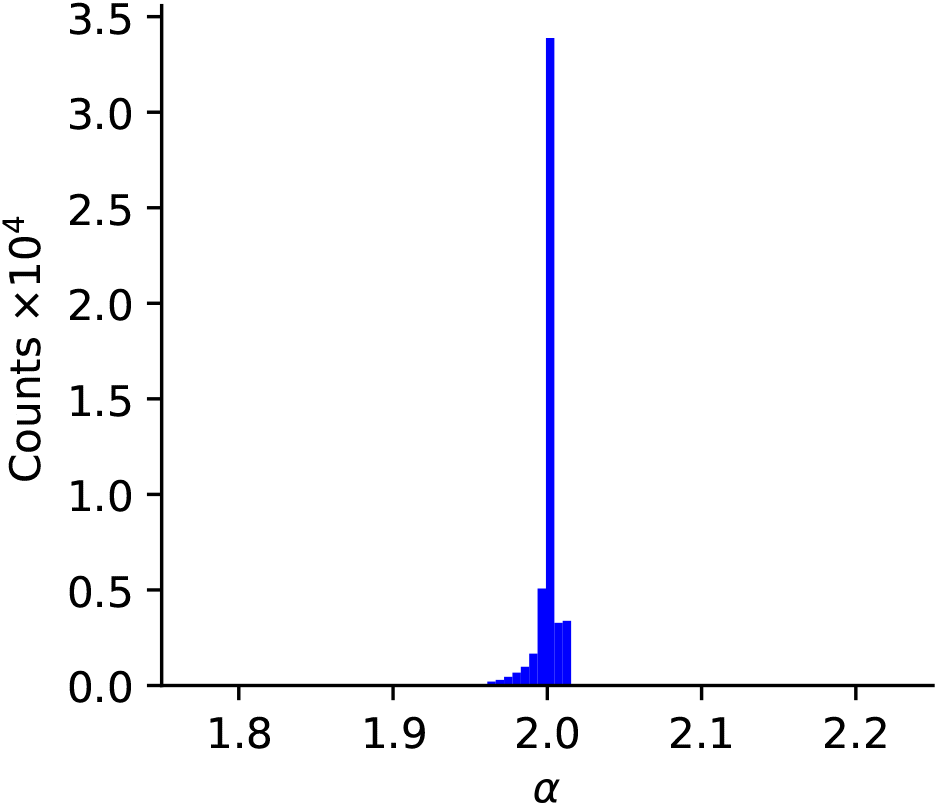
Approximating dwell time shape parameters of motors that maximize average run velocity at a given *k*_*h*_. Histogram of the shape parameters (*α*) for these motors show that they are all very close to two.

**Figure S3.**
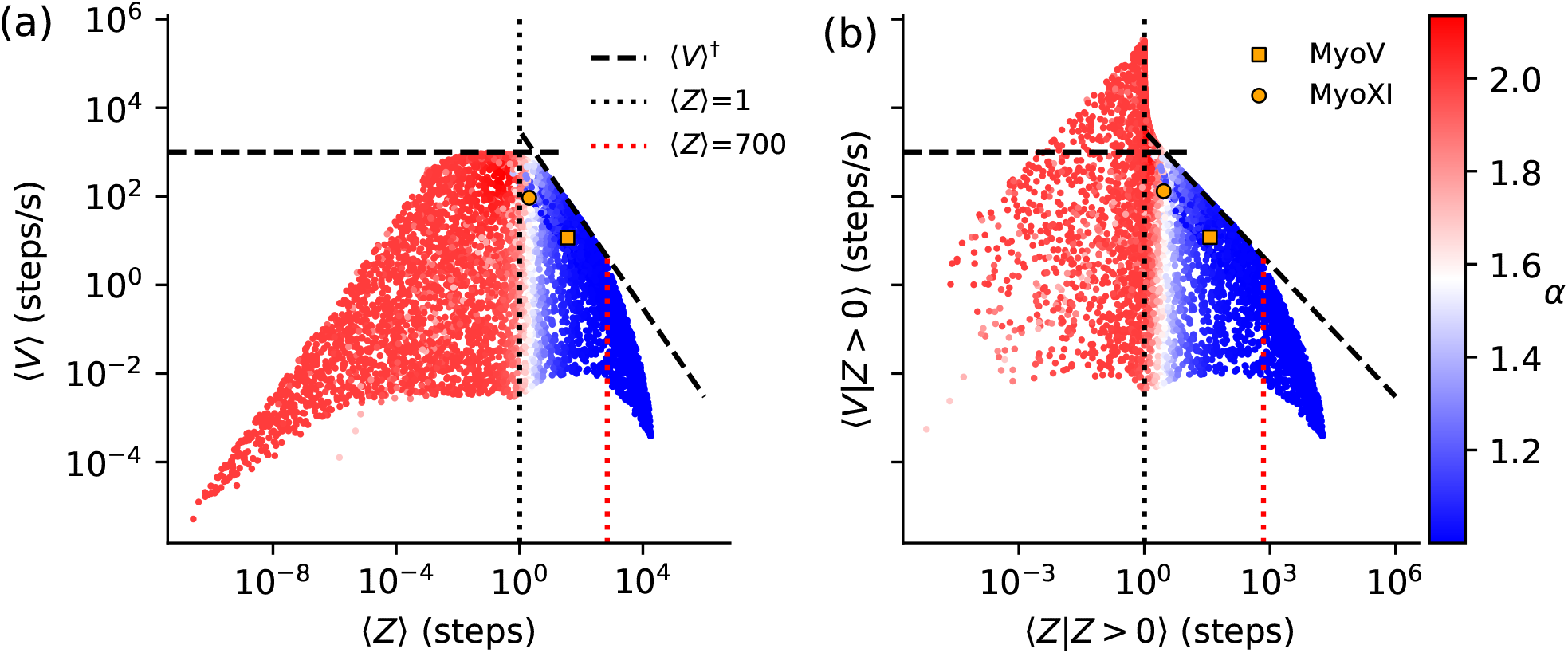
The average run velocity-distance Pareto frontier with imperfect gating. (a) Scatter plot of average run velocities (⟨*V*⟩) and run distances (⟨*Z*⟩) for 5000 motors with imperfect perfect gating, structural parameters like myosin-V^1^, and random values of *k*_*d*1_, *k*_*d*2_, and *k*_*h*_ drawn between 10^−2^ and 10^6^ on a log-uniform distribution. Points are color coded based on the value of their approximating parameter (*α*) and bounded by the Pareto frontier (⟨*V*⟩ ^†^. The vertical dotted line at ⟨*Z*⟩ = 1 separates the *α* = 1 (right, blue) and *α* = 2 (left, red) regions. The locations of myosin-V (MyoV,^1^) and *Nt*. myosin-XI (MyoXI, four parameter fitting results at 100 *µ*M ATP) are shown as an orange square and circle, respectively. (b) The same data as in panel (a) but with the average run velocities and run distances conditioned on productive runs (*Z* > 0). The vertical red dotted line at 700 steps indicates the upper limit of the loop used to calculate ⟨*V*⟩. The fall off of ⟨*V*⟩ to the right of 700 steps is a numerical artifact of this upper limit. Increasing it beyond 700 steps results in numerical instability.

**Figure S4.**
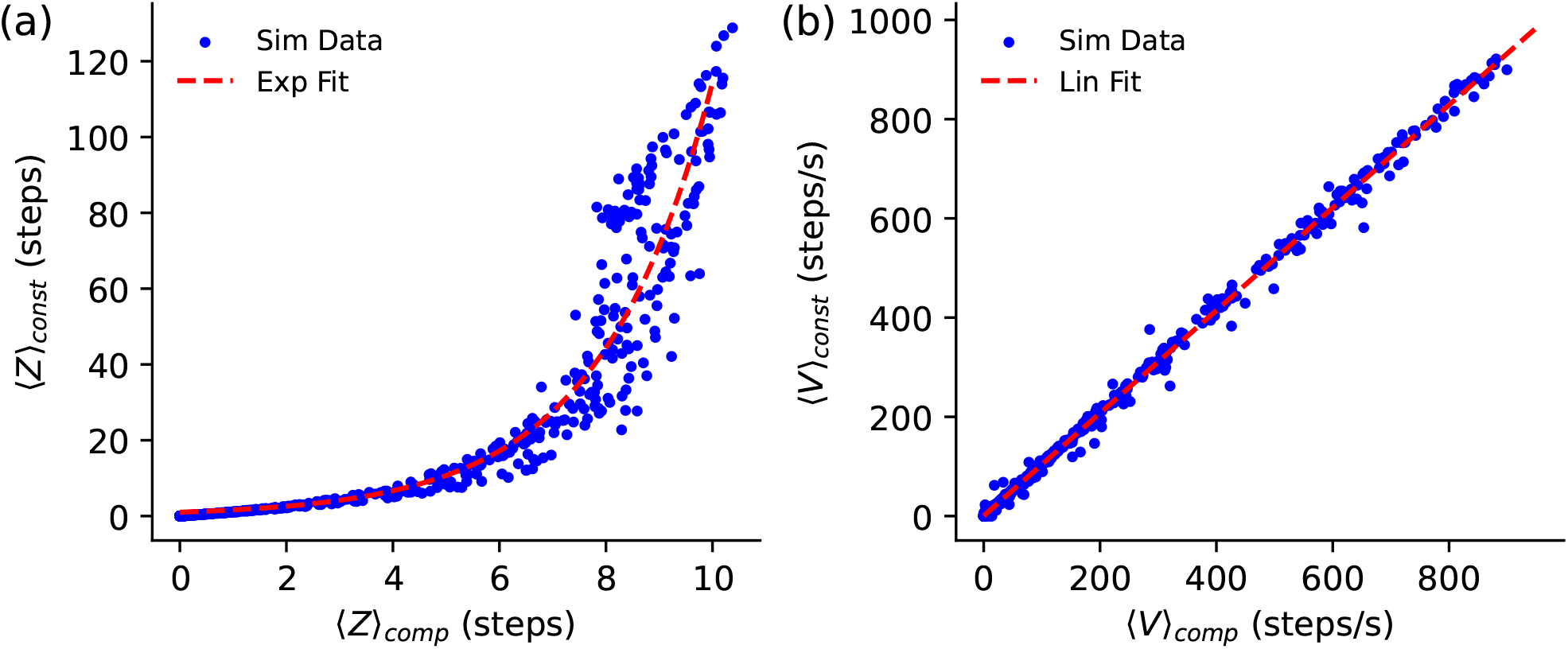
Average run distances and velocities from compliant and constant force optical traps are highly correlated. (a) The run distances show a non-linear correlation that is approximately exponential with rate 0.52. (b) The run velocities show linear correlation with slope of 1.04 and intercept 0.73, *R*^2^ = 0.998.

